# Author-Reviewer Homophily in Peer Review

**DOI:** 10.1101/400515

**Authors:** Dakota Murray, Kyle Siler, Vincent Larivière, Wei Mun Chan, Andrew M. Collings, Jennifer Raymond, Cassidy R. Sugimoto

## Abstract

The fairness of scholarly peer review has been challenged by evidence of disparities in publication outcomes based on author demographic characteristics. To assess this, we conducted an exploratory analysis of peer review outcomes of 23,876 initial submissions and 7,192 full submissions that were submitted to the biosciences journal *eLife* between 2012 and 2017. Women and authors from nations outside of North America and Europe were underrepresented both as gatekeepers (editors and peer reviewers) and authors. We found evidence of a homophilic relationship between the demographics of the gatekeepers and authors and the outcome of peer review; that is, there were higher rates of acceptance in the case of gender and country homophily. The acceptance rate for manuscripts with male last authors was seven percent, or 3.5 percentage points, greater than for female last authors (95% CI = [0.5, 6.4]); this gender inequity was greatest, at nine percent or about 4.8 percentage points (95% CI = [0.3, 9.1]), when the team of reviewers was all male; this difference was smaller and not significantly different for mixed-gender reviewer teams. Homogeny between countries of the gatekeeper and the corresponding author was also associated with higher acceptance rates for many countries. To test for the persistence of these effects after controlling for potentially confounding variables, we conducted a logistic regression including document and author metadata. Disparities in acceptance rates associated with gender and country of affiliation and the homophilic associations remained. We conclude with a discussion of mechanisms that could contribute to this effect, directions for future research, and policy implications. Code and anonymized data have been made available at https://github.com/murrayds/elife-analysis

**Author summary:** Peer review, the primary method by which scientific work is evaluated, is ideally a fair and equitable process in which scientific work is judged solely on its own merit. However, the integrity of peer review has been called into question based on evidence that outcomes often differ between male and female authors, and for authors in different countries. We investigated such disparities at the biosciences journal *eLife* by analyzing the demographics of authors and gatekeepers (editors and peer reviewers), and peer review outcomes of all submissions between 2012 and 2017. Outcomes were more favorable for male authors and those affiliated with institutions in North America and Europe; these groups were also over-represented among gatekeepers. There was evidence that peer review outcomes were influenced by *homophily* —a preference of gatekeepers for manuscripts from authors with shared characteristics. We discuss mechanisms that could contribute to this effect, directions for future research, and policy implications.

## Introduction

Peer review is foundational to the development, gatekeeping, and dissemination of research, while also underpinning the professional hierarchies of academia. Normatively, peer review is expected to follow the ideal of “universalism” [1], whereby scholarship is judged solely on its intellectual merit. However, confidence in the extent to which peer review promotes the best scholarship has been eroded by questions about whether social biases [2], based on or correlated with the demographic characteristics of the scholar, could also influence outcomes of peer review [3–5]. This challenge to the integrity of peer review has prompted increased interest in assessment of the disparities and potential influence of bias in their peer review processes.

Several terms are often conflated in the discussion of bias in peer review. We use the term *disparities* to refer to unequal composition between groups, *inequities* to characterize unequal outcomes, and *bias* to refer to the degree of impartiality in judgment or lack thereof. Disparities and inequities have been widely studied in scientific publishing, most notably with regard to gender and country of affiliation. Globally, women account for only about 30 percent of scientific authorships [6] and are underrepresented even when compared to their in the numbers in the scientific workforce [7, 8]. Underrepresentation of articles authored by women is most pronounced in the most prestigious and high-profile scientific journals [9–14]. Similar disparities are observed across countries, for which developed countries dominate the production of highly-cited publications [15, 16].

The under-representation of authors from certain groups may reflect differences in submission rates, or it may reflect differences in success rates during peer review (percent of submissions accepted), or both. Analyses of success rates have yielded mixed results in terms of the presence and magnitude of such inequities. Some analyses have found lower success rates for female-authored papers [17, 18] and grant applications [19, 20], while other studies have found no gender differences in review outcomes (for examples, see [21–25]). Inequities in journal success rates based on authors’ nationalities or country of affiliation have also been documented, with reports that authors from English-speaking and scientifically-advanced countries have higher success rates [26, 27]; however, other studies found no evidence that the language or country of affiliation of an author influences peer review outcomes [27–29]. These inconsistencies could be explained by several factors, such as the contextual characteristics of the studies (e.g., country, discipline) and variations in research design and sample size. Another possible explanation is that these gender and national disparities emerge from *bias* in peer review.

The possibility that bias contributes to inequities in scientific publishing and the nature of any such bias is highly controversial. Implicit bias—the macro-level social and cultural stereotypes that can subtly influence everyday interpersonal judgments and thereby produce and perpetuate status inequalities and hierarchies [30, 31]—has been suggested as a possible mechanism to explain differences in peer review outcomes based on socio-demographic and professional characteristics [3]. When faced with uncertainty—which is quite common in peer review—people often weight the social status and other ascriptive characteristics of others to help make decisions [32]. Hence, scholars are more likely to consider particularistic characteristics (e.g., gender, institutional prestige) of an author under conditions of uncertainty [33, 34], such as at the frontier of new scientific knowledge [35]. However, given the demographic stratification of scholars within institutions and across countries, it can be difficult to pinpoint the nature of a potential bias. For example, women are underrepresented in prestigious educational institutions [36–38], which conflates gender and prestige biases. These institutional differences can be compounded by gendered differences in age, professional seniority, research topic, and access to top mentors [39]. Another potential source of bias is what is dubbed cognitive particularism [40], whereby scholars harbor preferences for work and ideas similar to their own [41]. Evidence of this process has been reported in peer review in the reciprocity (i.e., correspondences between patterns of recommendations received by authors and patterns of recommendations given by reviewers in the same social group) between authors and reviewers of the same race and gender [42] (see also [43, 44]). Reciprocity can exacerbate or mitigate existing inequalities in science. If the work and ideas favored by gatekeepers are unevenly distributed across author demographics, this could be conducive to Matthew Effects [1], whereby scholars accrue accumulative advantages via *a priori* status privileges. Consistent with this, inclusion of more female reviewers was reported to attenuate biases that favored men in the awarding of RO1 grants at the National Institute of Health [18]. However, an inverse relationship was found in the evaluation of candidates for professorships [45] when female evaluators were present, male evaluators became less favorable toward female candidates. Thus the nature and potential impact of cognitive biases during peer review are multiple and complex.

Another challenge is to disentangle the contribution of bias during peer review from factors external to the review process that could influence success rates. For example, there are gendered differences in access to funding, domestic responsibilities, and cultural expectations of career preferences and ability [46, 47] that may adversely impact manuscript preparation and submission. On the other hand, women have been found to hold themselves to higher standards [48] and be less likely to compete [49], hence they may self-select a higher quality of work for submission to prestigious journals. At the country level, disparities in peer review outcomes could reflect structural factors related to a nation’s scientific investment [15, 50], publication incentives [51, 52], local challenges [53], and research culture [54], all of which could influence the actual and perceived quality of submissions from different nations. There are also several intersectional issues: there are, for example, differences in socio-demographic characteristics of the scientific workforce across countries—e.g., more women from some countries and disproportionately less professionally-senior women in others [6]. Because multiple factors external to the peer review process can influence peer review outcomes, unequal success rates for authors with particular characteristics do not necessarily reflect bias in the peer review process itself; conversely, equal success rates do not necessarily reflect a lack of bias.

Here, we assessed the extent to which bias contributes to gender and country disparities in peer review outcomes by analyzing the extent to which the magnitude of these disparities vary across different gender and country compositions of gatekeeper teams. In particular, we focused on the notion of homophily between the reviewers and authors. This analysis examined the outcomes of peer review at *eLife*—an open-access journal in the life and biomedical sciences. Peer review at *eLife* differs from other traditional forms of peer review used in the life sciences in that it is done through deliberation between reviewers (usually three in total) on an online platform. Previous studies have shown that deliberative scientific evaluation is influenced by social dynamics between evaluators [55, 56]. We examine how such social dynamics manifest in *eLife*’s deliberative peer review by assessing the extent to which the composition of reviewer teams correlates with peer review outcomes. Using all research papers (Research Articles, Short Reports, and Tools and Resources) submitted between 2012 and 2017 (n=23,876), we investigate the extent to which a relationship emerges between the gender and country of affiliation of authors (first, last, and corresponding) and gatekeepers (editors and invited peer reviewers), extending the approach used in previous work [2].

## Consultative peer review and *eLife*

Founded in 2012 by the Howard Hughes Medical Institute (United States), the Max Planck Society (Germany), and the Wellcome Trust (United Kingdom), *eLife* is an open-access journal that publishes research in the life and biomedical sciences. Manuscripts submitted to *eLife* progress through several stages. In the first stage, the manuscript is assessed by a Senior Editor, who may confer with one or more Reviewing Editors and decide whether to reject the manuscript or encourage the authors to provide a full submission. When a full manuscript is submitted, the Reviewing Editor recruits a small number of peer reviewers (typically two or three) to write reports on the manuscript. The Reviewing Editor is encouraged to serve as one of the peer reviewers. When all individual reports have been submitted, both the Reviewing Editor and peer reviewers discuss the manuscript and their reports using a private online discussion system hosted by *eLife*. At this stage the identities of the Reviewing Editor and peer reviewers are known to one another. If the consensus of this group is to reject the manuscript, all the reports are usually sent to the authors. If the consensus is that the manuscript requires revision, the Reviewing Editor and additional peer reviewers agree on the essential points that need to be addressed before the paper can be accepted. In this case, a decision letter outlining these points is sent to the authors (the original reports are not usually released in their entirety to the authors). When a manuscript is accepted, the decision letter and the authors’ response are published along with the manuscript. The name of the Reviewing Editor is also published. Peer reviewers can also choose to have their name published. This process has been referred to as consultative peer review (see [57, 58] for a more in-depth description of the *eLife* peer-review process).

## Data and methods

### Data

Metadata for research papers submitted to *eLife* between its inception in 2012 and mid-September, 2017 (n=23,876) were provided to us by *eLife* for analysis. As such, these data were considered a convenience sample. Submissions fell into three main categories: 20,948 Research Articles (87.7 percent), 2,186 Short Reports (9.2 percent), and 742 Tools and Resources (3.1 percent). Not included in this total were six Scientific Correspondence articles, which were excluded because they followed a distinct and separate review process. Each record potentially listed four submissions—an initial submission, full submission, and up to two revision submissions (though in some cases manuscripts remained in revision even after two revised submissions). Fig 1 depicts the flow of all 23,876 manuscripts through each review stage. The majority, 70.0 percent, of initial submissions for which a decision was made were rejected. Only 7,111 manuscripts were encouraged to submit a full submission. A total of 7,192 manuscripts were submitted as a full submission; this number was slightly larger than encouraged initial submissions due to appeals of initial decisions and other special circumstances. Most full submissions, 52.4 percent (n = 3,767), received a decision of revise, while 43.9 percent (n = 3,154) were rejected. A small number of full submissions (n = 54) were accepted without any revisions. On the date that data were collected (mid-September, 2017), a portion of initial submission (n = 147) and full submissions (n = 602) remained in various stages of processing and deliberation (without final decisions). On average, full submissions that were ultimately accepted underwent 1.23 revisions and, within our dataset, 3,426 full submissions were eventually accepted to be published. A breakdown of the number of revisions requested before a final decision was made, by gender and country of affiliation of the last author, is provided in S1 Fig. A portion of initial and full submissions (n = 619) appealed their decision, causing some movement from decisions of “Reject” to decisions of “Accept” or “Revise”; counts of appeals by the gender of author and gatekeepers is shown in S2 Fig.

**Fig 1.**
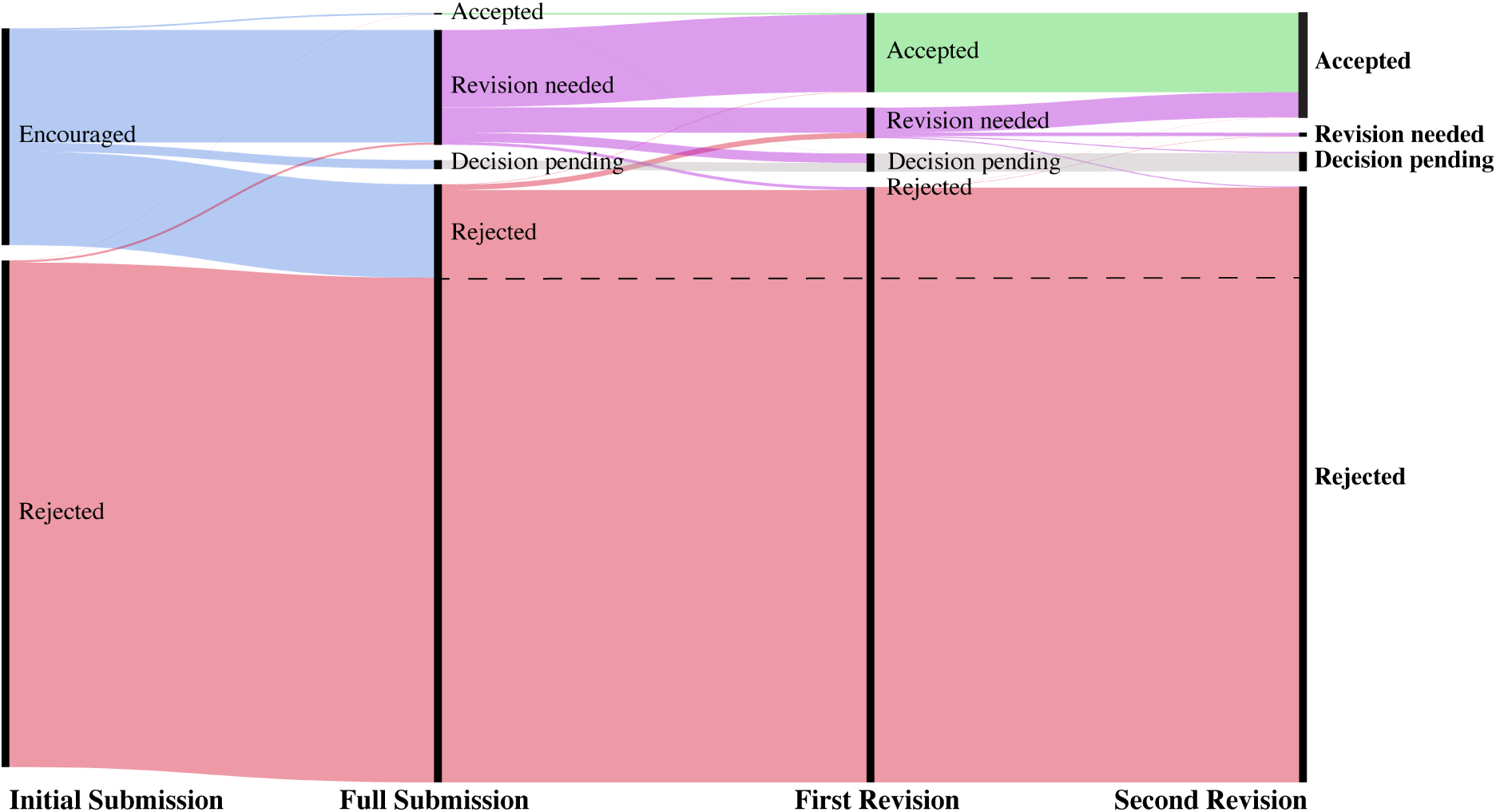
Flow of all papers through the *eLife* review process. Starting from the left, an initial submission is first given an initial decision of encourage or reject, and if encouraged, continues through the first full review and subsequent rounds of revision. “Encouraged”, “Accepted”, “Rejected” and “Revision needed” represent the decisions made by *eLife* editors and reviewers at each submission stage. A portion of manuscripts remained in various stages of processing at the time of data collection—these manuscripts were labeled as “Decision pending”. The status of manuscripts after the second revision is the final status that we consider in the present data. The dashed line delineates full submissions from rejected initial submissions.

The review process at *eLife* is highly selective, and became more selective over time. While only garnering 307 submissions in 2012, *eLife* accrued 8,061 submissions in 2016. Fig 2 shows that while the total count of manuscripts submitted to *eLife* has rapidly increased since the journal’s inception, the count of encouraged initial submissions and accepted full submissions has grown more slowly. The encourage rate (percentage of initial submissions encouraged to submit full manuscripts) was 44.6 percent in 2012, and dropped to 26.6 percent in 2016. The acceptance rate (the percentage of accepted full submissions) was 62.4 percent in 2012 and decreased to 53.0 percent in 2016. The overall acceptance rate (percentage of initial submissions eventually accepted) began at 27.0 percent in 2012 and decreased to 14.0 percent by 2016.

**Fig 2.**
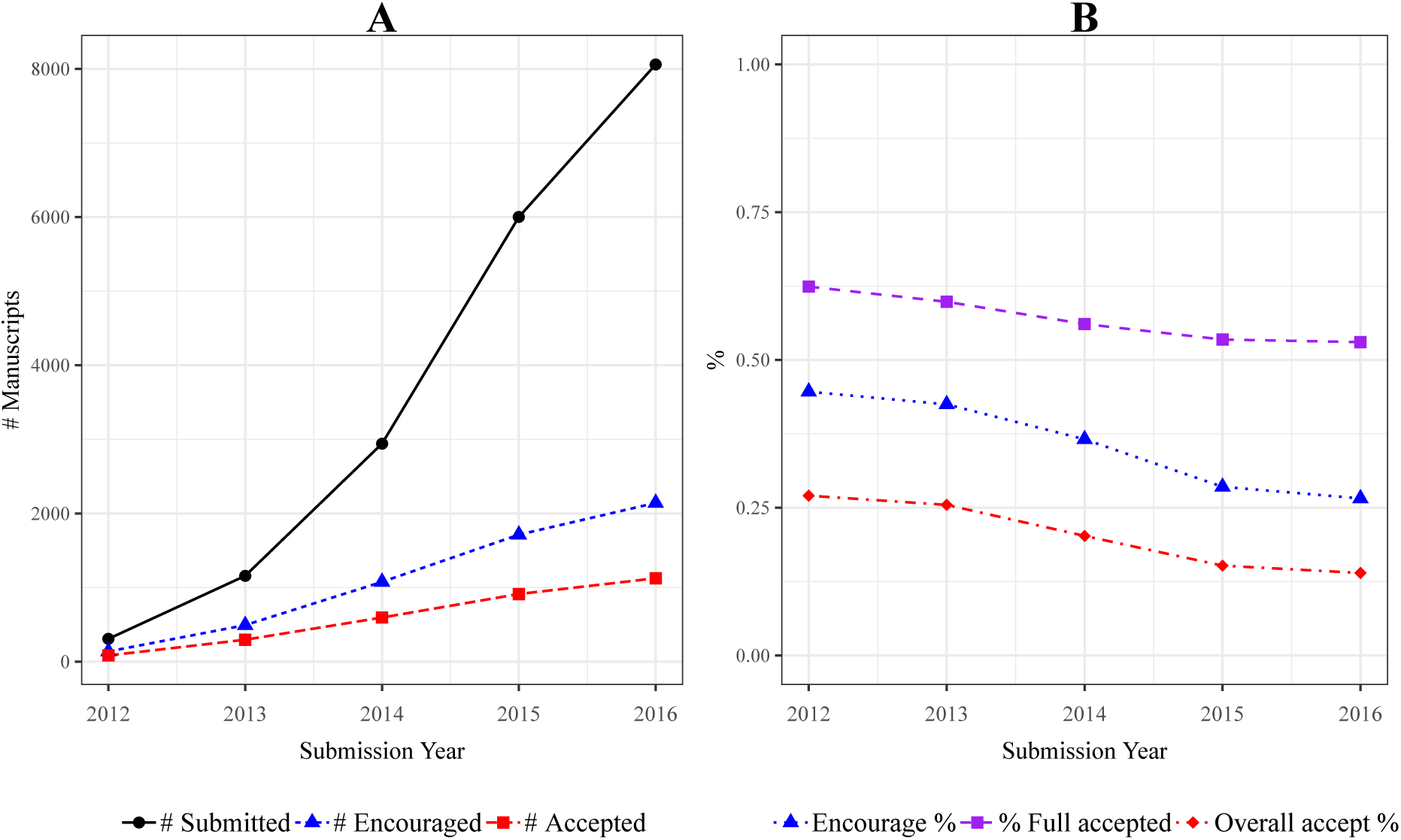
Submissions and selectivity of *eLife* over time. A: Yearly count of initial submissions, encouraged initial submissions, and accepted full submis-sions to *eLife* between 2012 and 2016; **B:** Rate of initial submissions encouraged (Encourage %), rate of full submissions accepted (% Full accepted) and rate of initial submissions accepted (Overall accept %) between 2012 and 2016. Submissions during the year of 2017 were excluded because we did not have sufficient data for full life-cycle of these manuscripts. Code to reproduce this figure can be found on the linked Github repository at the path figures/selectivity_over_time/selectivity_over_time.rmd.

In addition to authorship data, we obtained information about the gatekeepers involved in the processing of each submission. We defined gatekeepers as any Senior Editor or Reviewing Editor at *eLife* or invited peer reviewer involved in the review of at least one initial or full submission between 2012 and mid-September 2017. Gatekeepers at *eLife* often served in multiple roles; for example, acting as both a Reviewing Editor and peer reviewer on a given manuscript, or serving as a Senior Editor on one manuscript, but an invited peer review on another. In our sample, the Reviewing Editor was listed as a peer reviewer for 58.9 percent of full submissions. For initial submissions, we had data on only the corresponding author of the manuscript and the Senior Editor tasked with making the decision. For full submissions we had data on the corresponding author, first author, last author, Senior Editor, Reviewing Editor, and members of the team of invited peer reviewers. Data for each individual included their stated name, institutional affiliation, and country of affiliation. A small number of submissions were removed, such as those that had a first but no last author (reflecting compromised data record—even a single-authored manuscript should duplicate authors across all roles) and those that did not have a valid submission type. Country names were manually disambiguated (for example, by normalizing names such as “USA” to “United States” and “Viet Nam” to “Vietnam”). To simplify continent-level comparisons, we also excluded one submission for which the corresponding author listed their affiliation as Antarctica.

Full submissions included 6,669 distinct gatekeepers, 5,694 distinct corresponding authors, 6,691 distinct first authors, and 5,581 distinct last authors. Authors were also likely to appear on multiple manuscripts and may have held a different authorship role in each: whereas our data included 17,966 distinct combinations of author name and role, this number comprised only 12,059 distinct authors. For 26.5 percent of full submissions the corresponding author was also the first author, whereas for 71.2 percent of submissions the corresponding author was the last author. We did not have access to the full authorship list that included middle authors. Note that in the biosciences, the last author is typically the most senior researcher involved [59] and responsible for more conceptual work, whereas the first author is typically less senior and performs more of the scientific labor (such as lab work, analysis, etc.) to produce the study [60–62].

### Gender assignment

Gender variables for authors and gatekeepers were coded using an updated version of the algorithm developed in [6]. This algorithm used a combination of the first name and country of affiliation to assign each author’s gender on the basis of several universal and country-specific name-gender lists (e.g., United States Census). This list of names was complemented with an algorithm that searched Wikipedia for pronouns associated with names.

We validated this new list by applying it to a dataset of names with known gender. We used data collected from *RateMyProfessor.com*, a website containing anonymous student-submitted ratings and comments for professors, lecturers, and teachers for professors at universities in the United States, United Kingdom, and Canada. We limited the dataset to only individuals with at least five comments, and counted the total number of gendered pronouns that appeared in their text; if the total of one gendered-pronoun type was at least the square of the other, then we assigned the gender of the majority pronoun to the individual. To compare with pronoun-based assignment, we assigned gender using the previously detailed first-name based algorithm. In total, there were 384,127 profiles on *RateMyProfessor.com* that had at least five comments and for whom pronouns indicated a gender. Our first name-based algorithm assigned a gender of male or female to 91.26 percent of these profiles. The raw match-rate between these two assignments was 88.6 percent. Of those that were assigned a gender, our first name-based assignment matched the pronoun assignment in 97.1 percent of cases, and 90.3 percent of distinct first names. While *RateMyProfessor.com* and the authors submitting to *eLife* represent different populations (*RateMyProfessor.com* being biased towards teachers in the United States, United Kingdom, and Canada), the results of this validation provide some credibility to the first-name based gender assignment used here.

We also attempted to manually identify gender for all Senior Editors, Reviewing Editors, invited peer reviewers, and last authors for whom our algorithm did not assign a gender. We used Google to search for their name and institutional affiliation, and inspected the resulting photos and text in order to make a subjective judgment as to whether they were presenting as male or female.

Through the combination of manual efforts and our first-name based gender-assignment algorithm, we assigned a gender of male or female to 95.5 percent (n = 35,511) of the 37,198 name/role combinations that appeared in our dataset. 26.7 percent (n = 9,910) were assigned a gender of female, 68.8 percent (n = 25,601) were assigned a gender of male, while a gender assignment could be not assigned for the remaining 4.5 percent (n = 1,687). This gender distribution roughly matches the gender distribution observed globally across scientific publications [6]. A breakdown of these gender demographics by role can be found in S1 Table and S2 Table.

### Gender composition of reviewers

To assess the relationship between author-gatekeeper gender homogeny and review outcomes, we analyzed the gender composition of the gatekeepers and authors of full submissions. Each manuscript was assigned a reviewer composition category of *all-male*, *all-female*, *mixed*, or *uncertain*. Reviewer teams labeled *all-male* and *all-female* were teams for which we could identify a gender for every member, and for which all genders were identified as either male or female, respectively. Teams labeled as *mixed* were those teams for which we could identify a gender for at least two members, and which had at least one male and at least one female peer reviewer. Teams labeled as *uncertain* were those teams for which we could not assign a gender to every member and which were not mixed. A full submission was typically reviewed by two to three peer reviewers, which may or may not expicitely include the Reviewing Editor. However, the Reviewing Editor was always involved in the review process of a manuscript, and so we always considered the Reviewing Editor as a member of the reviewing team. Of 7,912 full submissions, a final decision of accept or reject was given for 6,590 during the dates analyzed; of these, 47.7 percent (n = 3,144) were reviewed by all-male teams, 1.4 percent (n = 93) by all-female teams, and 50.8 percent (n = 3,347) by mixed-gender teams; the remaining six manuscripts had reviewer teams classified as uncertain and were excluded from further analysis.

### Institutional Prestige

Institutional names for each author were added manually by *eLife* authors and were thus highly idiosyncratic. Many institutions appeared with multiple name variants (e.g., “UCLA”, “University of California, Los Angeles”, and “UC at Los Angeles”). In total, there were nearly 8,000 unique strings in the affiliation field. We performed several pre-processing steps on these names, including converting characters to lower case, removing stop words, removing punctuation, and reducing common words to abbreviated alternatives (e.g., “university” to “univ”). We used fuzzy-string matching with the Jaro-Winkler distance measure [63] to match institutional affiliations from *eLife* to institutional rankings in the 2016 *Times Higher Education World Rankings*. A match was established for 15,641 corresponding authors of initial submission (around 66 percent). Matches for last authors were higher: 5,118 (79 percent) were matched.

Institutions were classed into two levels of prestige: “top” institutions were those within the top 50 universities as ranked by the global *Times Higher Education* rankings. Institutions which ranked below the top 50, or which were otherwise unranked or which were not matched to a Times Higher Education ranking were labeled as “non-top”. One limitation of the Times Higher Education ranking as a proxy for institutional prestige is that these rankings cover only universities, excluding many prestigious research institutes. To mitigate this limitation, we mapped a small number of well-known and prestigious biomedical research institutes to the “top” category, including: The Max Plank Institutes, the National Institutes of Health, the UT Southwestern Medical Center, the Memorial Sloan Cancer Medical Center, the Ragon institutes, and the Broad Institute.

### Geographic distance

Latitude and longitude of country centroids were taken from Harvard WorldMap [64]; country names in the *eLife* and Harvard WorldMap dataset were manually disambiguated and then mapped to the country of affiliation listed for each author from *eLife* (for example, ”Czech Republic” from the *eLife* data was mapped to ”Czech Rep.” in the Harvard WorldMap data). For each initial submission, we calculated the geographic distance between the centroids of the countries of the corresponding author and Senior Editor; we call this the *corresponding author-editor geographic distance*. For each full submission, we calculated the sum of the geographic distances between the centroid of the last author’s country and the country of each of the reviewers. All distances were calculated in thousands of kilometers; we call this the *last author-reviewers geographic distance*.

### Analysis

We conducted a series of *χ*^2^ tests of equal proportion as well as several logistic regression models in order to assess the likelihood that an initial submission is encouraged and that a full submission is accepted, as a function of author and gatekeeper characteristics. We supply p-values and confidence intervals as a tool for interpretation; we generally maintain the convention of 0.05 as the threshold for statistical significance, though we also report and interpret values just outside of this range. When visualizing proportions, 95% confidence intervals are calculated using the definition 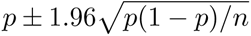, where *p* is the proportion and *n* is the number of observations in the group. When conducting *χ*^2^ tests comparing groups based on gender, we excluded submissions for which no gender could be identified. When conducting tests for gender and country homogeny, we report 95% interval confidence intervals of their difference in proportion—we do not report confidence intervals for tests involving more than two groups. Odds ratios and associated 95% confidence intervals are reported for logistic regression models. Data processing, statistical testing, and visualization was performed using R version 3.4.2 and RStudio version 1.1.383.

Having conducted an exploratory analysis of gender and country inequities in peer review with this univariate approach, we built a series of logistic regression models to investigate whether these differences could be explained by other factors. In each model, we used the submission’s outcome as the response variable, whether that be encouragement (for initial submissions) or acceptance (for full submissions). For both initial and full submissions, we added control variables for the year of submission (measured from 0 to 5, representing 2012 to 2017, accordingly), the type of the submission (Research Article, Short Report, or Tools and Resources), and the institutional prestige of the author (top vs non-top). For full submissions, we also controlled for the gender of the first author. Mirroring the univariate analysis, we constructed two sets of models. The first set of models investigates the extent of peer review inequities based on author characteristics. We considered predictor variables for the gender and continent of affiliation of the corresponding author (for initial submissions), and the last author (for full submissions). For the second set of models, we investigated whether these inequities differed based on gender or country homogeny between the author and the reviewer or editor. In addition to variables from the first model, we considered several approaches to capture the effect of gender-homogeny between the author and reviewers on peer review inequity (see below). We also included variables for the corresponding author-editor geographic distance (for initial submissions), and last author-reviewers geographic distance (for full submission), and a dummy variable indicating whether this distance was zero; these variables serve as proxies for the degree of country homogeny between the author and the editor or reviewers. There were a small number of Senior Editors in our data—in order to protect their identity we did not include their gender or specific continent of affiliation in any models; we maintained a variable for corresponding author-editor geographic distance.

Several approaches were considered for modeling the relationship between equity in peer review and the composition of the reviewer team using logistic regression. Approaches such as modelling equity using simple interaction terms or with a two-model approach were also considered but were ultimately excluded due to methodological and interpretive constraints (see S1 Text and S2 Text for more discussion of these models and their results). A third approach modelled equity across groups as a categorical variable consisting of all six combinations of last author gender (male, female) and reviewer team composition (all-male, all-female, mixed); This approach provides a more interpretable means of testing the extent to which gender equity in success rates was related to the interaction between author and reviewer team demographics, and was the focus of our analysis.

## Results

### Gatekeeper representation

We first analyzed whether the gender and countries of affiliation of the population of gatekeepers at *eLife* was similar to that of the authors of initial and full submissions. The population of gatekeepers comprised primarily of invited peer reviewers, as there were far fewer Senior and Reviewing Editors. A gender and country breakdown by gatekeeper type has been provided in S2 Table, and S3 Table.

Fig 3 illustrates the gender and country demographics of authors and gatekeepers. The population of gatekeepers at *eLife* was largely male. Only 21.6 percent (n = 1,440) of gatekeepers were identified as female, compared with 26.6 percent (n = 4,857) of corresponding authors (includes authors of initial submissions), 33.9 percent (n = 2,272) of first authors, and 24.0 percent (n = 1,341) of last authors. For initial submissions, we observed a strong difference between the gender composition of gatekeepers and corresponding authors, *χ*^2^(df= 1*, n* = 17, 119) = 453.9*, p ≤* 0.00001. The same held for full submissions, with a strong difference for first authorship, *χ*^2^(df= 1*, n* = 6, 153) = 844.4*, p ≤* 0.0001; corresponding authorship, *χ*^2^(df= 1*, n* = 6, 647) = 330.04*, p ≤* 0.0001; and last authorship, *χ*^2^(df= 1*, n* = 5, 292) = 17.7*, p ≤* 0.00003. Thus, the gender proportions of gatekeepers at *eLife* was male-skewed in comparison to the authorship profile.

**Fig 3.**
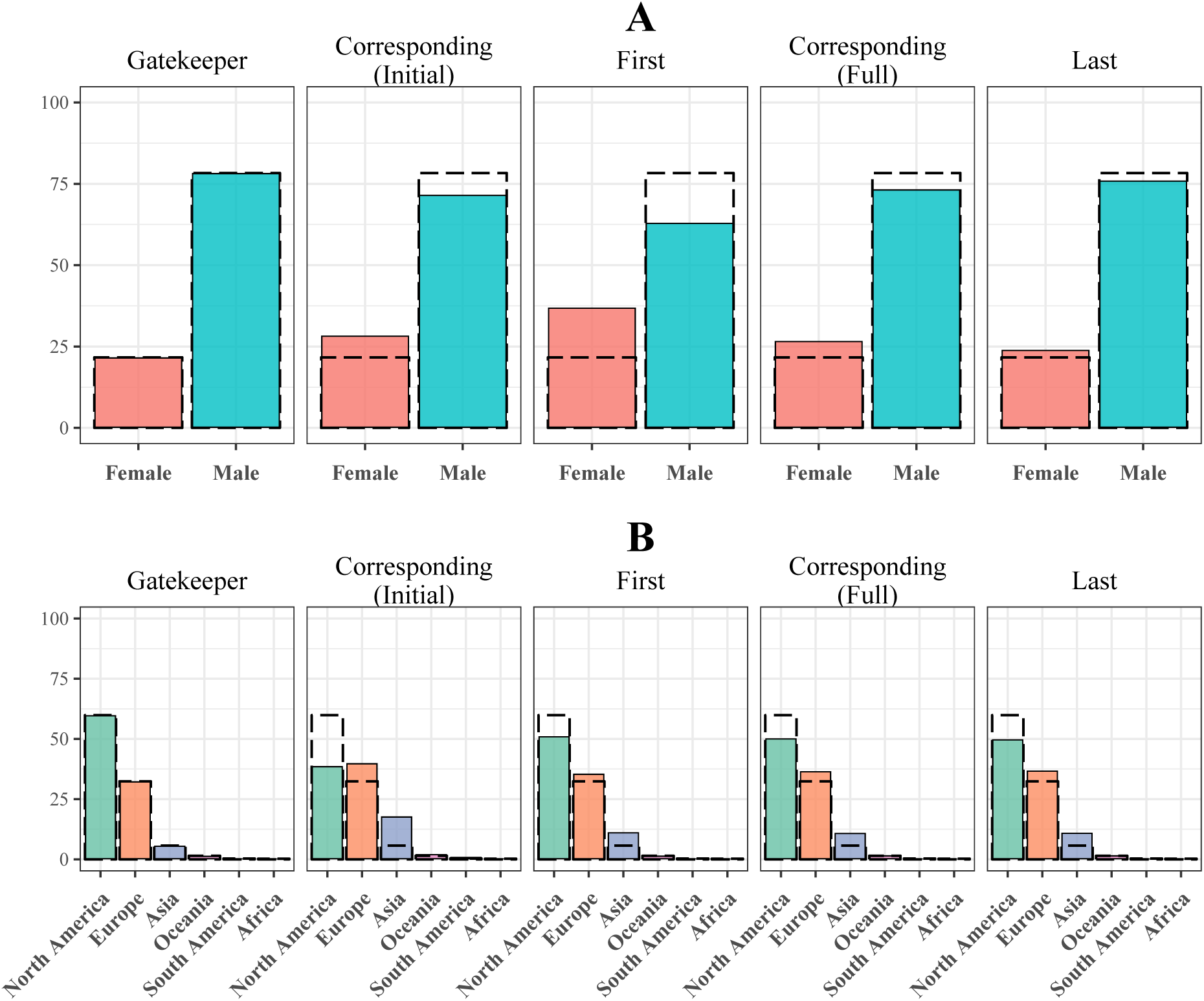
Gender and country of affiliation demographics of authors and gatekeepers at *eLife*. **A:** proportion of identified men and women in the populations of distinct gatekeepers (Senior Editors, Reviewing Editors, and peer reviewers) and of the populations of distinct corresponding authors on initial submissions, and first, corresponding and last authors on full submissions; percentages exclude those for whom no gender was identified. **B:** proportion of people with countries of affiliation within each of six continents in the population of distinct gatekeepers, and for the population of distinct corresponding, first, and last authors. Black dashed lines overlaid on authorship graphs indicate the proportion of gatekeepers within that gendered or continental category. Values used in this graph can be found in S1 Table and S4 Table. Code to reproduce this figure can be found on the linked Github repository at the path figures/gatekeeper_representation/gatekeeper_representation.rmd.

The population of gatekeepers at *eLife* was heavily dominated by those from North America, who constituted 59.9 percent (n = 3,992) of the total. Gatekeepers from Europe were the next most represented, constituting 32.4 percent (n = 2,162), followed by Asia with 5.7 percent (n = 378). Individuals from South America, Africa, and Oceania each made up less than two percent of the population of gatekeepers. As with gender, we observed differences between the country composition of gatekeepers and that of the authors. Gatekeepers from North America were over-represented whereas gatekeepers from Asia and Europe were under-represented for all authorship roles. For initial submissions, there was a significant difference in the distribution of corresponding authors compared to gatekeepers *χ*^2^(df= 5*, n* = 18, 195) = 6738.5*, p ≤* 0.00001. The same held for full submissions, with a significant difference for first authors, *χ*^2^(df= 5*, n* = 6, 674) = 473.3*, p ≤* 0.00001, corresponding authors, *χ*^2^(df= 5*, n* = 6, 669) = 330.04*, p ≤* 0.00001, and last authors *χ*^2^(df= 5*, n* = 5, 595) = 417.2*, p ≤* 0.0001. The international representation of gatekeepers was most similar to first and last authorship (full submissions), and least similar to corresponding authorship (initial submissions) due to country-level differences in acceptance rates (see Fig 4). We also note that the geographic composition of submissions to *eLife* has changed over time, attracting more submissions from authors in Asia in later years of analysis (see S4 Fig).

**Fig 4.**
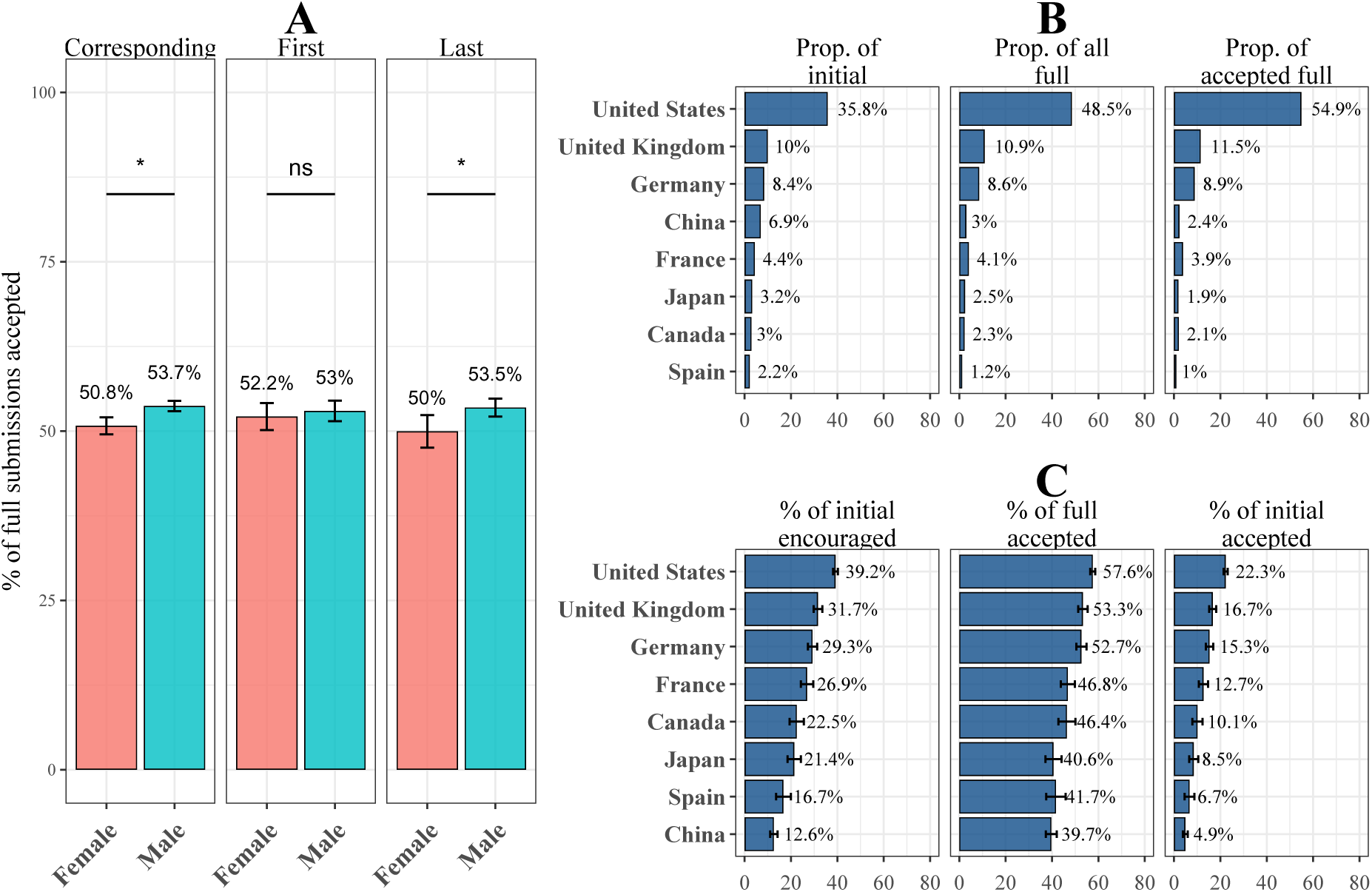
Peer review success rates by gender and country of authors. A: Percentage of full submissions that were accepted, shown by the gender of the corresponding author, first author, and last author. Authors whose gender was unknown were excluded from analysis. See S3 Fig for an extension of this figure including submission rates, encourage rates, and overall acceptance races. Error bars indicate 95% confidence intervals of the proportion of accepted full submissions. Asterisks indicate significance level of *χ*^2^ tests of independence of frequency of acceptance by gender; “***” = *p* < 0.001; “**” = *p* < 0.01; “*” = *p* < 0.05; “-” = *p* < 0.1; “ns” = *p ≥* 0.1. **B:** Proportion of all initial submissions, full submissions, and accepted full submissions by the country of affiliation of the corresponding author for the top eight most prolific countries in terms of initial submissions. **C:** Encourage rate of initial submissions, acceptance rate of full submissions, and acceptance rate of initial submissions by country of affiliation of the corresponding author for the top eight more prolific countries in terms of initial submissions. Error bars indicate 95% confidence intervals for each percentage. This same graph with the top 16 most prolific nations can be found in S7 Fig. Code to reproduce this figure can be found on the linked Github repository at the path figures/author_outcomes/submission_outcome_by_gender.rmd.

### Peer review success rates by author gender, country of affiliation

Male authorship dominated *eLife* submissions: men accounted for 76.9 percent (n = 5,529) of gender-identified last authorships and 70.7 percent (n = 5,083) of gender-identified corresponding authorships of full submissions (see S3 Fig). First authorship of full submissions was closest to gender parity, although still skewed towards male authorship at 58.1 percent (n = 4,179).

We observed a gender inequity favoring men in the outcomes of each stage of the review processes. The percentage of initial submissions encouraged was 2.1 percentage points higher for male corresponding authors—30.83 to 28.75 percent, *χ*^2^(df= 1*, n* = 22, 319) = 8.95, 95% CI = [0.7, 3.4]*, p* = 0.0028 (see S3 Fig). Likewise, the percentage of full submissions accepted was higher for male corresponding authors—53.7 to 50.8 percent *χ*^2^(df= 1*, n* = 6, 188) = 3.95, 95% CI = [0.03, 5.8]*, p* = 0.047. The gender inequity at each stage of the review process yielded higher overall acceptance rates (the percentage of initial submissions eventually accepted) for male corresponding authors (15.6 percent) compared with female corresponding authors (13.8 percent), *χ*^2^(df= 1*, n* = 21, 670) = 10.96, 95% CI = [0.8, 2.9]*, p* = 0.0009 for a male:female success ratio of 1.13 to 1.

Gender disparity was only apparent in the senior authorship roles. Fig 4.A shows the gendered acceptance rates of full submissions for corresponding, first and last authors. There was little to no relationship between the gender of the first author and the percentage of full submissions accepted, *χ*^2^(df= 1*, n* = 5, 971) = 0.34, 95% CI = [*−*1.8, 3.5]*, p* = 0.56. There was, however, a significant gender inequity in full submission outcomes for last authors, as also observed for corresponding authors—the acceptance rate of full submissions was 3.5 percentage points higher for male as compared to female last authors—53.5 to 50.0 percent, *χ*^2^(df= 1*, n* = 6, 505) = 5.55, 95% CI = [0.5, 6.4]*, p* = 0.018.

Fig 4.B shows the proportion of manuscripts submitted, encouraged, and accepted to *eLife* from corresponding authors originating from the eight most prolific countries (in terms of initial submissions). Manuscripts with corresponding authors from these eight countries accounted for a total of 73.9 percent of all initial submissions, 81.2 percent of all full submissions, and 86.5 percent of all accepted publications. Many countries were underrepresented in full and accepted manuscripts compared to their submissions. For example, whereas papers with Chinese corresponding authors accounted for 6.9 percent of initial submissions, they comprised only 3.0 percent of full and 2.4 percent of accepted submissions. The only countries that were over-represented—making up a greater portion of full and accepted submissions than expected given their initial submissions—were the United States, United Kingdom, and Germany. In particular, corresponding authors from the United States made up 35.8 percent of initial submissions, yet constituted 48.5 percent of full submissions and the majority (54.9 percent) of accepted submissions.

Each stage of review contributed to the disparity of country representation between initial, full, and accepted submissions, with manuscripts from the United States, United Kingdom, and Germany more often encouraged as initial submissions and accepted as full submissions. Fig 4.C shows that initial submissions with a corresponding author from the United States were the most likely to be encouraged (39.2 percent), followed by the United Kingdom (31.7 percent) and Germany (29.3 percent). By contrast, manuscripts with corresponding authors from Japan, Spain, and China were comparatively less likely to be encouraged (21.4, 16.7, and 12.6 percent, respectively). These differences narrowed somewhat for full submissions: the acceptance rate for full submissions with corresponding authors from the U.S. was the highest (57.6 percent), though more similar to other countries, such as the United Kingdom, Germany, and France than for encourage rates.

There were gendered differences in submissions by country of affiliation (S5 Fig), but there were insufficient data to test whether gender and country of affiliation interacted to affect the probability of acceptance.

The gender and country inequities evident in the univariate analyses were subsequently affirmed using a logistic regression model that controlled for a number of potential confounds (see Fig 5. We modeled peer review outcomes based on not only the gender and affiliated continent of the corresponding author (initial submissions) and last author (full submissions), but also the prestige of the author’s institution, the year in which the manuscript was submitted, and the submission type (Research Article, Short Report, or Tools and Resources). For full submissions, we also controlled for the gender of the first author. The results of this regression for initial and full submissions are shown in Fig 5.

**Fig 5.**
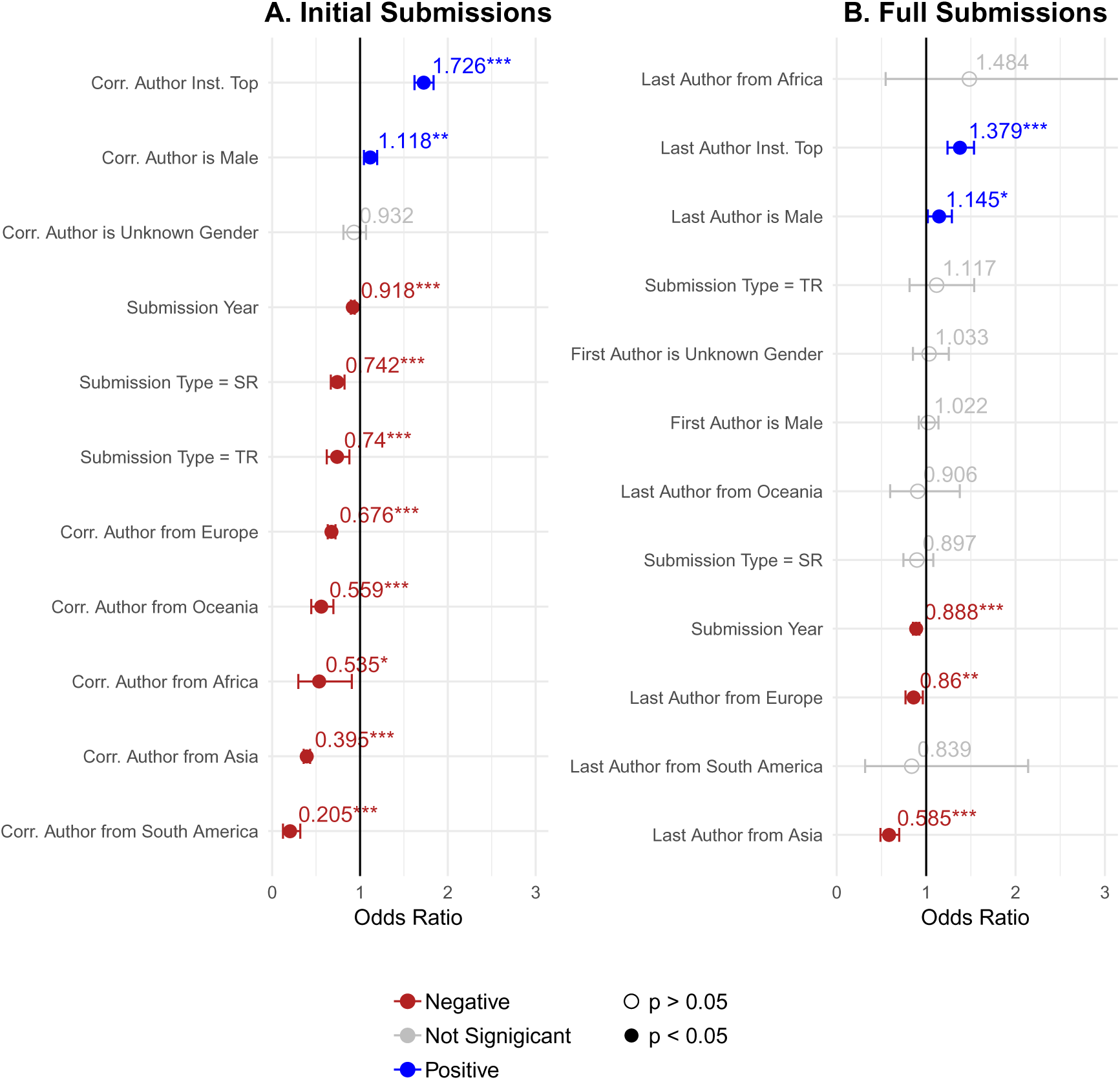
Modelling success rates of initial and full submissions based on author characteristics. A: Estimates of a logistic regression model of initial submissions using whether the submission was encouraged as the response variable, and available information on the corresponding author as predictors. **B:** Estimates of a logistic regression model of full submissions using whether the submission was accepted as the response variable, and available information about the first and last authors as predictors. For both initial and full submissions, control variables included author’s institutional prestige, the year of submission, and the submission type. For full submissions, there is also a control variable for the gender of the first author. For continent of affiliation, we held “North America” as the reference level. For submission type, “RA” (research article) was used as the reference level; the submission type “SR” means “Short Reports”, and “TR” means “Tools and Resources”. Blue, red, and grey points indicate positive, negative, and non-significant effects, respectively. The numbers above each point label the size of the effect, as an odds ratio. Bars extending from either side of each point indicate 95% confidence intervals. Asterisks next to each label indicate significance level: “***” = *p ≤* 0.001; “**” = *p ≤* 0.01; “*” = *p ≤* 0.05; otherwise, *p >* 0.05. Some confidence intervals are cropped; a table detailing full effects are included in S6 Table and S7 Table. Code to reproduce this figure can be found on the linked Github repository at the path figures/regression_analysis/regression_analysis_simple.rmd.

For both initial and full submissions, the prestige of the author’s institution was the strongest predictor of a positive peer review outcome (initial: *β* = 1.726, 95% CI = [1.663, 1.789]*, p ≤* 0.0001; full: *β* = 1.379, 95% CI = [1.272, 1.486]*, p ≤* 0.0001). A more recent year of submission was associated with a lower odds of acceptance, (initial: *β* = 0.918, 95% CI = [0.894, 0.942]*, p ≤* 0.0001; full: *β* = 0.888, 95% CI = [0.847, 0.929]*, p ≤* 0.0001), reflecting the increasing selectivity of *eLife* (see Fig 2). Compared to Research Articles, both Short Reports, (*β* = 0.742, 95% CI = [0.638, 0.847]*, p ≤* 0.0001), and Tools and Resources (*β* = 0.740, 95% CI = [0.567, 0.913]*, p ≤* 0.0001) were less likely to have a positive review outcome at the initial submission stage.

Even when controlling for these variables, there were still inequities by the gender and country of affiliation of the author, affirming trends illustrated in Fig 4. Initial submissions with a male corresponding author were associated with a 1.12 times increased odds of being encouraged (95% CI = [1.048, 1.182]*, p* = 0.0014), and full submissions with a male last author with a 1.14 times increased odds of being accepted (95% CI = [1.03, 1.26]*, p* = 0.025), compared to submissions with female corresponding or last authors. In contrast, for the first author position, there was no significant difference in outcomes by gender. The logistic regression also provided evidence for the presence of geographic inequities, with lower odds of success for submissions with authors outside of North America. Compared to submissions with a corresponding author from North America, an initial submission with a corresponding author from Europe was 0.68 times as likely to be encouraged (95% CI = [0.3236, 0.783]*, p ≤* 0.0001), and a corresponding author from Oceania was 0.56 times as likely to be encouraged (95% CI = [0.34, 0.78]*, p ≤* 0.0001), followed by corresponding authors from Africa (*β* = 0.53, 95% CI = [*−*0.18, 1.088]*, p* = 0.027), Asia (*β* = 0.40, 95% CI = [0.30, 0.49]*, p ≤* 0.0001), and South America (*β* = 0.21, 95% CI = [*−*0.269, 0.679]*, p ≤* 0.0001). Geographic disparities were also present, although less pronounced for full submissions, with significantly lower odds of acceptance for submissions with a last author from Europe (*β* = 0.86, 95% CI = [0.75, 0.97]*, p* = 0.008) or Asia (*β* = 0.59, 95% CI = [0.41, 0.76]*, p ≤* 0.0001) compared with North America.

### Peer Review Outcomes by Author-Gatekeeper Homogeny

The higher acceptance rates for male authors manifested largely from instances when the reviewer team was all male (Fig 6). When all reviewers were male, the acceptance rate of full submissions was about 4.7 percentage points higher for male compared to female last authors (*χ*^2^ = 4.48(df= 1*, n* = 3, 110), 95% CI = [0.3, 9.1], *p* = 0.034) and about 4.4 points higher for male compared to female corresponding authors (S6 Fig; *χ*^2^(df= 1*, n* = 2, 974) = 3.97, 95% CI = [0.1, 8.7]*p* = 0.046). For mixed-gender reviewer teams, the disparity in author success rates by gender was smaller and not statistically-significant. All-female reviewer teams were too rare to draw firm conclusions (only 81 of 6,509 processed full submissions), but in the few cases of all-female reviewer teams, there was a higher acceptance rate for female last, corresponding, and first authors that did not reach statistical significance. There was no significant relationship between first authorship gender and acceptance rates, regardless of the gender composition of the reviewer team. In sum, greater parity in outcomes was observed when gatekeeper teams contained both men and women. Notably, the acceptance rate for female authors was not lower for all-male reviewer teams compared with mixed reviewer teams, rather the gender disparity arose from a higher acceptance rate for submissions from male authors when they were reviewed by a team of all-male reviewers. We refer to this favoring by reviewers of authors sharing their same gender as *homophily*.

**Fig 6.**
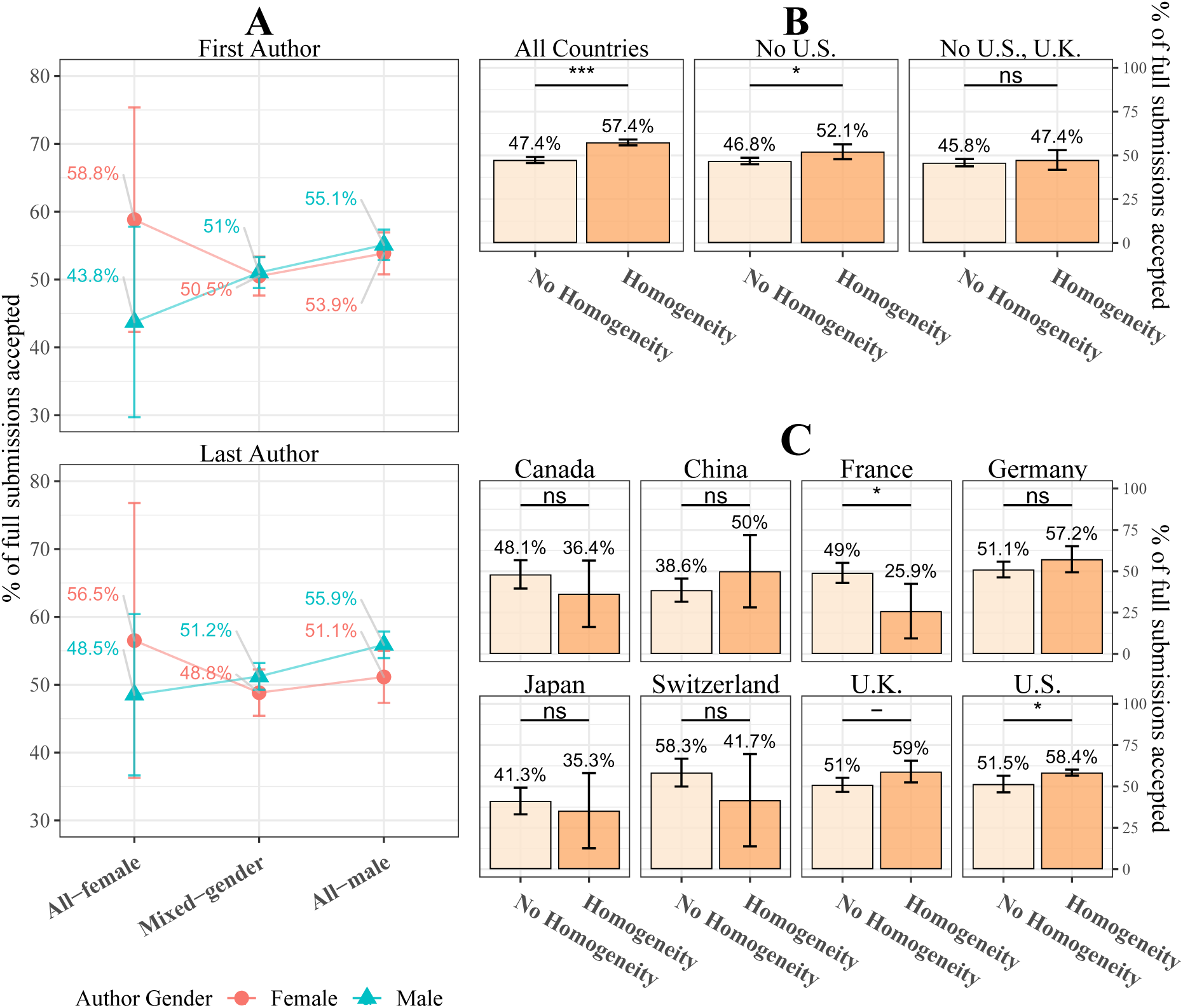
Relationship between author-reviewer homogeny and peer review outcomes. A: Percentage of full submissions that were accepted by gender of the first author (top) and last author (bottom), partitioned by the gender composition of the peer reviewers. The y-axis has been cropped between 30 percent and 80 percent in order to draw attention to the relevant effect. See S6 Fig for more information. **B:** Peer review outcome by presence of country homogeny (last author from the same country as at least one reviewer) for all submissions (left), excluding submissions from the United States (middle) and excluding submissions from the United States and the United Kingdom, the two countries with the highest acceptance rates (right). **C:** Acceptance rate of full submissions by country homogeny, shown for individual countries. Shown are the top eight most prolific countries in terms of number of initial submissions. For all panels: vertical error bars indicate 95% percentile confidence intervals. Values at the base of each bar indicate the number of observations within each group. Asterisks indicate significance level of *χ*^2^ tests of independence comparing frequency of accepted full submissions between presence and absence of homogeny and within each country. “***” = *p* < 0.001; “**” = *p* < 0.01; “*” = *p* < 0.05; “-”= *p* < 0.1; “ns” = *p ≥* 0.1. Code to reproduce this figure can be found on the linked Github repository at the path figures/gatekeeper_author_outcomes/gatekeeper_author_outcomes.rmd.

Homophily was also evident in the relationship between peer review outcomes and the presence of country homogeny between the last author and reviewer. We defined last author-reviewer country homogeny as a condition for which at least one member of the reviewer team (Reviewing Editor and peer reviewers) listed the same country of affiliation as the last author. We only considered the country of affiliation of the last author, since it was the same as that of the first and corresponding author for 98.4 and 94.9 percent of full submissions, respectively. Outside of the United States, the presence of country homogeny during review was rare. Whereas 88.4 percent of full submissions with last authors from the U.S. were reviewed by at least one gatekeeper from their country, country homogeny was present for only 29.3 percent of full submissions with last authors from the United Kingdom and 26.2 percent of those with a last author from Germany. The incidence of reviewer homogeny fell sharply for Japan and China which had geographic homogeny for only 10.3 and 9.9 percent of full submissions, respectively. More extensive details on the rate of author/reviewer homogeny for each country can be found in S5 Table.

Last author-reviewer country homogeny tended to result in the favoring of submissions from authors of the same country as the reviewer. We first pooled together all authors from all countries (n = 6,508 for which there was a full submission and a final decision), and found that the presence of homogeny during review was associated with a 10.0 percentage point higher acceptance rate, (Fig 6.B; *χ*^2^(1*, n* = 6, 508) = 65.07, 95% CI = [7.58, 12.47]*, p ≤* 0.00001). However, most cases of homogeny occurred for authors from the United States, so this result could potentially reflect the higher acceptance rate for these authors (see Fig 4), rather than homophily overall. Therefore we repeated the test, excluding all full submissions with last authors from the United States, and we again found a significant, though statistically less confident homophilic effect, *χ*^2^(df= 1*, n* = 3, 236) = 4.74, 95% CI = [0.52, 10.1]*, p* = 0.029. We repeated this procedure again, excluding authors from both the United States and United Kingdom, (the two nations with the highest acceptance rates, see 4), and we identified no homophilic effect, *χ*^2^(df= 1*, n* = 1, 920) = 0.016, 95% CI = [*−*4.6, 7.7]*p* = 0.65. Thus, the effects of last-author reviewer country-homophily were largely driven by the United States and United Kingdom.

For authors from outside the United States, not only was the presence of author-reviewer country homogeny rare, but the tendency for a homophilic effect on peer review outcome appeared to vary, depending on the country. Fig 6.C shows acceptance rates for last authors affiliated within the eight most prolific nations submitting to *eLife*. For the United States, presence of homogeny was associated with a 6.9 percentage point higher likelihood of acceptance compared to no homogeny *χ*^2^(df= 1*, n* = 3, 270) = 6.25, 95% CI = [1.4, 12.4]*, p* = 0.0124. Similarly, papers from the United kingdom were 8.0 percentage points more likely to be accepted if there was last author-reviewer homogeny *χ*^2^(df= 1*, n* = 739) = 3.65, 95% CI = [*−*0.1, 16.2]*, p* = 0.056. In contrast, submissions with last authors from France were 23 percentage points *less* likely to be accepted if there was country homogeny *χ*^2^(df= 1*, n* = 204) = 4.34, 95% CI = [*−*42.8*, −*3.4]*, p* = 0.037. There was a similar, though non-significant effect for Canada and Switzerland (also French-speaking countries). Due to the rarity of country homogeny outside of the U.S., more data are needed to draw firm conclusions on a per-country basis.

To further assess the contribution of author-reviewer homogeny to inequity in peer review outcomes, we extended the logistic regression approach shown in Fig 5. For full submissions, we compared two logistic regression models, one that considered author-reviewer geographic homogeny but only main effects of reviewer team gender composition (Fig 7.A) and one that included terms to model the effects of author-reviewer geographic and gender homogeny (Fig 7.B). To model the extent to which gender equity differed based on the gender composition of the reviewer team, we modelled interactions using a variable combining factor levels for last author gender and reviewer team composition (Fig 7.B). To model the degree of country homogeny between the author and the author and the reviewers, we included in the model the last author-reviewers geographic distance, defined as the sum of the geographic distance between the centroids of the last author’s country, and the country of all of the peer reviewers. All distances were calculated in thousands of kilometers; for example, the geographic distance between the United States and Denmark is 7.53 thousands of kilometers. We included a dummy variable indicating whether the distance was zero. A similar analysis was performed to assess the effect of author-editor homogeny on the outcomes of initial submissions (S8 Table); this excludes any analysis of homophily between the author and Senior Editor in order to protect the identity of the small number of Senior Editors.

**Fig 7.**
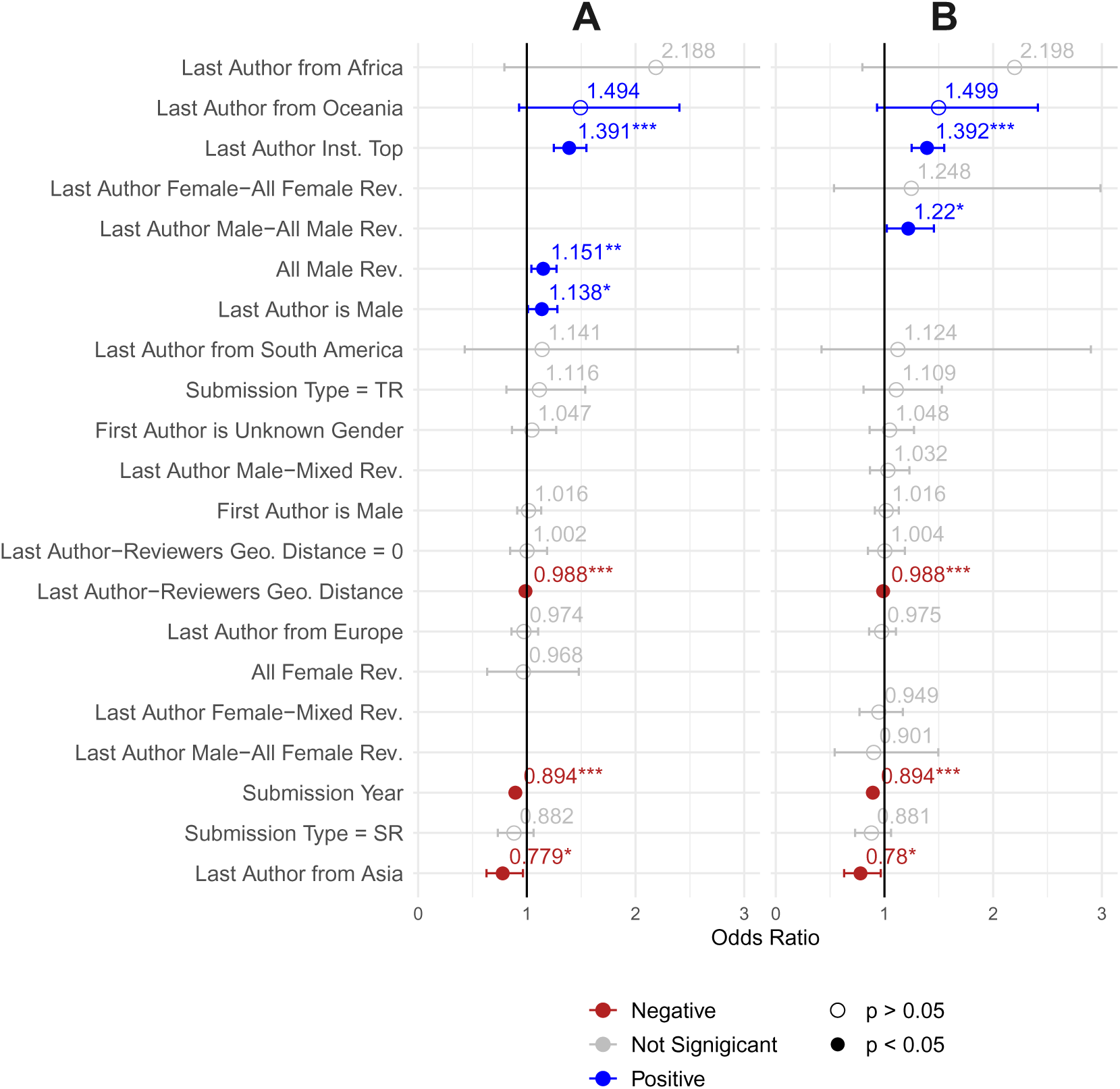
Modelling success of full submissions with author-reviewer homogeny. Estimates of logistic regression models of full submissions using whether the submission was accepted as the response variable. **A:** Includes as predictors the demographic and geographic characteristics of last author and gatekeepers, along with an indicator or the level of last author-reviewer geographic homogeny. **B:** Includes all predictors as in **A** but with the last author gender and reviewer gender composition combined into a single, six-level categorical variable. Control variables for both panels include author’s institutional prestige, year of submission, submission type, and gender of the first author. For continent of affiliation, “North America” was used as the reference level. For submission type, “RA” (research article) was used as the reference level; the submission type “SR” means “Short Reports”, and “TR” means “Tools and Resources”. For the combination variable of last author gender and reviewer team gender composition, we held “last author female—all rev. male” as the reference level. Blue and red points indicate positive and negative effects, respectively. The numbers above each point are the size of the effect as an odds ratio. Bars extending from either side of each point indicate 95% confidence intervals. Asterisks above each label indicate significance level: “***” = *p* < 0.001; “**” = *p* < 0.01; “*” = *p* < 0.05; otherwise, *p >* 0.05. Some confidence intervals are cropped; a table detailing full effects is included in S9 Table. Code to reproduce this figure can be found on the linked Github repository at the path figures/regression_analysis/regression_analysis_interaction.rmd.

Fig 7.A shows that there were similar main effects of author gender and country, in terms of direction and magnitude, as in Fig 5.B. Even after controlling for reviewer team composition, a full submission with a male last author was 1.14 times more likely to be accepted than a submission with a female last author (95% CI = [1.020, 1.256]*, p* = 0.032). In addition, there were inequities based on author continent of affiliation, although smaller than in Fig 5.B. Affiliation within Asia was associated with a 0.779 times reduced odds of acceptance compared to North America (95% CI = [0.565, 0.992]*, p* = 0.022)—a smaller effect size than the 0.585 times reduced odds observed in Fig 5.B. Submissions with a last author from Oceania were associated with a 1.494 times increased odds of acceptance compared to North America, though with wide confidence intervals (95% CI = [1.020, 1.968]*, p* = 0.097]); this diverges from the non-significant negative effect observed in Fig 5. The effect of control variables—submission year, submission type, author institutional prestige, and first author gender—were also similar to those in Fig 5.

The extended model in Fig 7.A revealed a main effect of reviewer team gender composition. Compared to mixed-gender reviewer teams, submissions reviewed by all-male reviewers were 1.15 times more likely to be accepted (95% CI = [1.051, 1.252]*, p* = 0.0059); there was no significant difference between all-female and mixed-gender teams. In addition, this model revealed an influence of author-reviewer geographic homogeny. Every 1000km of last author-reviewer distance was associated with a 0.988 times lower odds of acceptance (95% CI = [0.982, 0.994]*, p ≤* 0.0001). This negative effect of last author-reviewers geographic distance provides additional evidence for the observations from Fig 6—that homogeny between the author and reviewers was associated with a greater odds of acceptance, even when controlling for the continent of affiliation of the author and other characteristics of the author and submission. A last author-reviewers geographic distance of zero (indicating that all reviewers were from the same country as the corresponding author) was not associated with a strong effect beyond that predicted by distance.

Finally, we modelled interactions between last author gender and reviewer-team composition by combining them into a single categorical variable containing all six combinations of factor levels (Fig 7.B). Full submissions with a male last author and which were reviewed by a team of all-male reviewers was associated with a 1.22 times higher odds of being accepted than a full submission with a female last author that was reviewed by an all male team (95% CI = [1.044, 1.40]*, p* = 0.027). No significant differences were observed for other combinations of author gender and reviewer gender composition. The absolute difference in parameter estimates between male and female authors among mixed-gender teams (0.084) was less than half that of all-male reviewer teams (0.198), suggesting greater equity among submissions reviewed by mixed-gender teams than by all-male teams. Taken together, these findings suggest that gender inequity in peer review outcomes tended to be smaller for mixed-gender reviewer teams, even controlling for many potentially confounding factors. These results provide evidence affirming observations from the univariate analysis in Fig 6.

## Discussion

We identified inequities in peer review outcomes at *eLife*, based on the gender and country of affiliation of the senior (last and corresponding) authors. Acceptance rates were higher for male than female last authors. In addition, submissions from developed countries with high scientific capacities tended to have higher success rates than others. These inequities in peer review outcomes could be attributed, at least in part, to a favorable interaction between gatekeeper and author demographics under the conditions of gender or country homogeny; we describe this favoring as *homophily*, a preference based on shared characteristics. Gatekeepers were more likely to recommend a manuscript for acceptance if they shared demographic characteristics with the authors, demonstrating homophily. In particular, manuscripts with male (last or corresponding) authors had a significantly higher chance of acceptance than female (last or corresponding) authors when reviewed by an all male review team. Similarly, manuscripts tended to be accepted more often when at least one of the reviewers was from the same country as the corresponding author (for initial submissions) or the last author (for full submissions), though there may be exceptions on a per-country basis (such as France and Canada). We followed our univariate analysis with a regression analysis, and observed evidence that these inequities persisted even when controlling for potentially confounding variables. The differential outcomes on the basis of author-reviewer homogeny is consistent with the notion that peer review at *eLife* is influenced by some form of bias—be it implicit bias [3, 17], geographic or linguistic bias [26, 65, 66], or cognitive particularism [40]. Specifically, homophilic interaction suggests that peer review outcomes may sometimes be associated with factors other than the intrinsic quality of a manuscript, such as the composition of the review team.

The opportunity for homophilous interactions is determined by the demographics of the gatekeeper pool, and the demographics of the gatekeepers differed significantly from those of the authors, even for last authors, who tend to be more senior [59–62]. The underrepresentation of women at *eLife* mirrors global trends—women comprise a minority of total authorships, yet constitute an even smaller proportion of gatekeepers across many domains [14, 67–74]. Similarly, gatekeepers at *eLife* were less geographically diverse than their authorship, reflecting the general underrepresentation of the “global south” in leadership positions of international journals [75].

The demographics of the reviewer pool made certain authors more likely to benefit from homophily in the review process than others. Male lead authors had a nearly 50 percent chance of being reviewed by a homophilous (all-male), rather than a mixed-gender team. In contrast, because all-female reviewer panels were so rare (accounting for only 81 of 6,509 full submission decisions), female authors were highly unlikely to benefit from homophily in the review process. Similarly, U.S. authors were much more likely than not (see S5 Table) to be reviewed by a panel with at least one reviewer from their country. However, the opposite was true for authors from other countries. Fewer opportunities for such homophily may result in a disadvantage for scientists from smaller and less scientifically prolific countries.

Increasing representation of women and scientists from a more diverse set of nations among *eLife’s* editor may lead to more diverse pool of peer reviewers and reviewing editors and a more equitable peer review process. Editors often invite peer reviewers from their own professional networks, networks that likely reflect the characteristics of the editor [76–78]; this can lead to editors, who tend to be men [14, 67–74] and from scientifically advanced countries [75] to invite peer reviewers who are demographically similar to themselves [44, 79, 80], inadvertently excluding certain groups from the gatekeeping process. Accordingly, we found that male Reviewing Editors at *eLife* were less likely to create mixed-gender teams of gatekeepers than female Reviewing Editors (see S8 Fig). We observed a similar effect based on the country of affiliation of the Reviewing Editor and invited peer reviewers (see S9 Fig). Moreover, in S11 Table we conducted a regression analysis considering only the gender of the Reviewing Editor, rather than the composition of the reviewer team; we found similar homophilous relationships as in Fig 7, suggesting the importance of the reviewing editor to the peer review process at *eLife*.

The size of disparities we observed in peer review outcomes may seem modest; however these small disparities accumulate through each stage of the review process (initial submission, full submission, revisions). These cumulative effects yield an overall acceptance rate (the rate at which initial submissions were eventually accepted) for male and female corresponding authors of 15.6 and 13.8 percent respectively; in other words, manuscripts submitted to *eLife* with male corresponding were published at a rate 1.13 times the rate of those with female corresponding authors. Similarly, initial submissions by corresponding authors from China were accepted at only 22.0 percent the rate of manuscripts submitted by corresponding authors from the United States (with overall acceptance rates of 4.9 and 22.3 percent, respectively). Success in peer review is vital for a researcher’s career because successful publication strengthens their professional reputation and makes it easier to attract funding, students, postdocs, and hence further publications. Even small advantages can compound over time and result in pronounced inequalities in science [81–84].

Our finding that the gender of the last authors was associated with a significant difference in the rate at which full submissions were accepted at *eLife* stands in contrast with a number of previous studies of journal peer review that reported no significant difference in outcomes of papers submitted by male and female authors [85–87], or differences in reviewer’s evaluations based on the author’s apparent gender [88]. This discrepancy may be explained in part by *eLife’s* unique context, policies, or the relative selectivity of *eLife* compared to journals where previous studies found gender equity. In addition, our results point to a key feature of study design that may account for some of the differences across studies: the consideration of multiple authorship roles. This is especially important for the life sciences, for which authorship order is strongly associated with contribution [61, 62, 89]. Whereas our study examined the gender of the first, last, and corresponding authors, most previous studies have focused on the gender of the first author (e.g., [2, 90]) or of the corresponding author (e.g., [22, 91]). Consistent with previous studies, we observed no strong relationship between first author gender and review outcomes at *eLife*. Only when considering lead authorship roles—last authorship, and to a lesser extent, corresponding author, did we observe such an effect. Our results may be better compared with studies of grant peer review, where leadership roles are more explicitly defined, and many studies have identified significant disparities in outcomes favoring men [18, 92–95], although many other studies have found no evidence of gender disparity [21, 23, 24, 96–98]. Given that science has grown increasingly collaborative and that average authorship per paper has expanded [99, 100], future studies of disparities would benefit from explicitly accounting for multiple authorship roles and signaling among various leadership positions on the byline [59, 101].

The relationship we found between the gender and country of affiliation of gatekeepers and peer review outcomes also stands in contrast to the findings from a number of previous studies. Studies of gatekeeper country of affiliation have found no difference in peer review outcomes based on the country of affiliation or country of affiliation of the reviewer [104, 106], though there is little research on the correspondence between author and reviewer gender. One study identified a homophilous relationship between female reviewers and female authors, [102]. However, most previous analyses found only procedural differences based on the gender of the gatekeeper [22, 87, 88, 103] and identified no difference in outcomes based on the interaction of author and gatekeeper gender in journal submissions [87, 104, 105] or grant review [23]. One past study examined the interaction between U.S. and non-U.S. authors and gatekeepers, but found an effect opposite to what we observed, such that U.S. reviewers tended to rate submissions of U.S. authors more harshly than those of non-U.S. authors [43]. Our results also contrast with the study most similar to our own, which found no evidence of bias related to gender, and only modest evidence of bias related to geographic region [2]. These discrepancies may result from our analysis of multiple author roles rather than considering only the characteristics of the first author. Alternatively, they may result from the unique nature of *eLife*’s consultative peer review; the direct communication between peer reviewers compared to traditional peer review may render the social characteristics of reviewers more influential.

### Limitations

There are limitations of our methodology that must be considered. First, we have no objective measure of the intrinsic quality of manuscripts. Therefore, it is not clear which review condition (homophilic or non-homophilic) more closely approximates the ideal of merit-based peer review outcomes. Second, measuring the relationship between reviewer and author demographics on peer review outcomes cannot readily detect biases that are shared by all reviewers/gatekeepers (e.g., if all reviewers, regardless of gender, favored manuscripts from male authors); hence, our approach could underestimate the influence of bias. Third, our analysis is observational, so we cannot establish causal relationships between success rates and authors or gatekeeper demographics—there remain potential confounding factors that we were unable to control for in the present analysis, such as the gender distribution of submission by country (see S5 Fig). Along these lines, the reliance on statistical tests with arbitrary significance thresholds may provide misleading results (see [107]), or obfuscate statistically weak but potentially important relationships. Fourth, our gender-assignment algorithm is only a proxy for author gender and varies in reliability by continent.

Further studies will be required to determine the extent to which the effects we observed generalize to other peer review contexts. Specific policies at *eLife*, such as their consultative peer review process, may contribute to the effects we observed. Other characteristics of *eLife* may also be relevant, including its level of prestige [13], and its disciplinary specialization in the biological sciences, whose culture may differ from other scientific and academic disciplines. It is necessary to determine the extent to which the findings here are particularistic or generalizable; it may also be useful in identifying explanatory models. Future work is necessary to confirm and expand upon our findings, assess the extent to which they can be generalized, establish causal relationships, and mitigate the effects of these methodological limitations. To aid in this effort, we have made as much as possible of the data and analysis publicly available at: https://github.com/murrayds/elife-analysis.

## Conclusion and recommendations

Many factors can contribute to gender, country, and other inequities in scientific publishing [47, 50, 108–111], which can affect the quantity and perceived quality of submitted manuscripts. However, these structural factors do not readily account for the observed effect of gatekeeper-author demographic homogeny associated with peer review outcomes at *eLife*; rather, relationships between the personal characteristics of the authors and gatekeepers are likely to play some role in peer review outcomes.

Our results suggest that it is not only the form of peer review that matters, but also the composition of reviewers. Homophilous preferences in evaluation are a potential mechanism underpinning the Matthew Effect [1] in academia. This effect entrenches privileged groups while potentially limiting diversity, which could hinder scientific advances, since diversity may lead to better working groups [112] and promote high-quality science [113, 114]. Increasing gender and international representation among scientific gatekeepers may improve fairness and equity in peer review outcomes and accelerate scientific progress. However, this must be carefully balanced to avoid overburdening scholars from minority groups with disproportionate service obligations.

Although some journals and publishers, such as *eLife* and Frontiers Media, have begun providing peer review data to researchers (see [44, 115]), data on equity in peer review outcomes is currently available only for a small fraction of journals and funders. While many journals collect these data internally, they are not usually standardized or shared publicly. One group, *PEERE*, authored a protocol for open sharing of peer review data [116, 117], though this protocol is recent, and the extent to which it will be adopted remains uncertain. Watchdog groups, such as BiaswatchNeuro, are now tracking and posting the representation of women authors in some journals. To both provide better benchmarks and to incentivize better practices, journals should make analyses on author and reviewer demographics publicly available. These data include, but would not be limited to, characteristics such as gender, race, sexual orientation, seniority, and institution and country of affiliation. It is likely that privacy concerns and issues relating to confidentiality will limit the full availability of the data; but analyses that are sensitive to the vulnerabilities of smaller populations should be conducted and made available as benchmarking data. As these data become increasingly available, systematic reviews can be useful in identifying general patterns across disciplines and countries.

Some high-profile journals have experimented with implementing double-blind peer review as a potential solution to inequities in publishing, including Nature [118] and *eNeuro* [12], though in some cases with low uptake [119]. Our findings of homophilic effects may suggest that single-blind review is not the optimal form of peer review; however, our study did not directly test whether homophily persists in the case of double blind review. If homophily is removed in double-blind review, it would reinforce the interpretation of bias; if it is maintained, it would suggest other underlying attributes of the manuscript that may be contributing to homophilic effects. Double-blind peer review is viewed positively by the scientific community [120, 121], and some studies have found evidence that double-blind review mitigates inequities that favor famous authors, elite institutions [85, 122, 123], and those from high-income and English-speaking nations [28].

There may be a tension, however, in attempting to further double blind peer review while other aspects of the scientific system become more open. More than 20 percent of *eLife* papers that go out for review, for example, are already available as preprints, which complicates the possibility of truly blind review. To a lesser extent, several statements required for the responsible conduct of research—e.g., conflicts of interest, funding statements, and other ethical declarations—would require altered administrative treatment to implement double blind review. Other options involve making peer review more open—one recent study showed evidence that more open peer review did not compromise the integrity or logistics of the process, so long as reviewers could maintain anonymity [124].

Other alternatives to traditional peer review have also been proposed, including study pre-registration, consultative peer review, and hybrid processes (eg: [58, 125–129]), as well as alternative forms of dissemination, such as preprint servers (e.g., arXiv, bioRxiv) which have in recent years grown increasingly popular [130]. Currently, there is little empirical evidence to determine whether these formats constitute more equitable alternatives [3]. In addition, some journals are analyzing the demographics of their published authorship and editorial staff in order to identify key problem areas, focus initiatives, and track progress in achieving diversity goals [14, 79, 86]. More work should be done to study and understand the issues facing peer review and scientific gatekeeping in all its forms and to promote fair, efficient, and meritocratic scientific cultures and practices. Editorial bodies should craft policies and implement practices to mitigate disparities in peer review; they should also continue to be innovative and reflective about their practices to ensure that papers are accepted on scientific merit, rather than particularistic characteristics of the authors.

## Supporting information

**S1 Text Modelling homogeny using main effects with interaction term.** We used logistic regression to model the degree to which gender equity in peer review outcomes differed based on the composition of the reviewer team in order to verify the inequity observed in Fig 6. Fig 7.A demonstrates that last author gender inequity persisted even when controlling for the gender composition of the reviewer team, but did not address the degree to which this equity manifests in submissions reviewed by all-male vs. mixed-gender reviewer teams. Given that there is no established method of addressing this question, we considered several approaches. The first approach modelled the interaction between last author gender and the gender-composition of the reviewer team (see S9 Table, column 2), however this approach proved difficult to interpret: adding the interaction term appeared to suppress the main effects of last author gender and reviewer team composition observed in Fig 7.A, though the corresponding ANOVA table demonstrated these effects to still account for a significant amount of deviance (see S11 Table). There were no significant interaction term, conflicting with Fig 6; main effects are often made less interpretable by the addition of interaction terms. A low sample size across interaction groups further complicates interpretation. Moreover, this approach modelled individual-level interactions between the author and reviewer composition on a per-submission basis, not differences in group-level estimates of inequity.

**S2 Text Modelling homogeny using separately trained models.** S9 Table, columns 3 and 4 shows the results of two logistic regression models of percentage of full submissions accepted, constructed as in fig 7.A, but each calculated using only full submissions reviewed by either all-male or mixed-gender reviewer teams. In the all-male model, a male last author was associated with a 1.23 times increased odds of acceptance (95% CI = [1.05, 1.41]*, p* = 0.027) compared to a female last author; in contrast, a smaller non-significant effect was observed between male and female last authors in the model containing only mixed-gender reviewer teams. This approach shows a larger positive effect favoring male last authors under the condition of all-male teams than for mixed-reviewer teams, affirming results of models in Fig 7.B, but this approach has several limitations that favor the approach from Fig 7. The confidence intervals for the effect of the regression for submissions reviewed by mixed reviewer teams are wide, making precise comparisons difficult. Interpretation of S9 Table is further complicated by possible population differences between groups as well as the different amount of data used to fit each model, n=3,090 for the all-male reviewer model and n = 3,280 for the mixed-gender reviewer model.

**S1 Fig.**
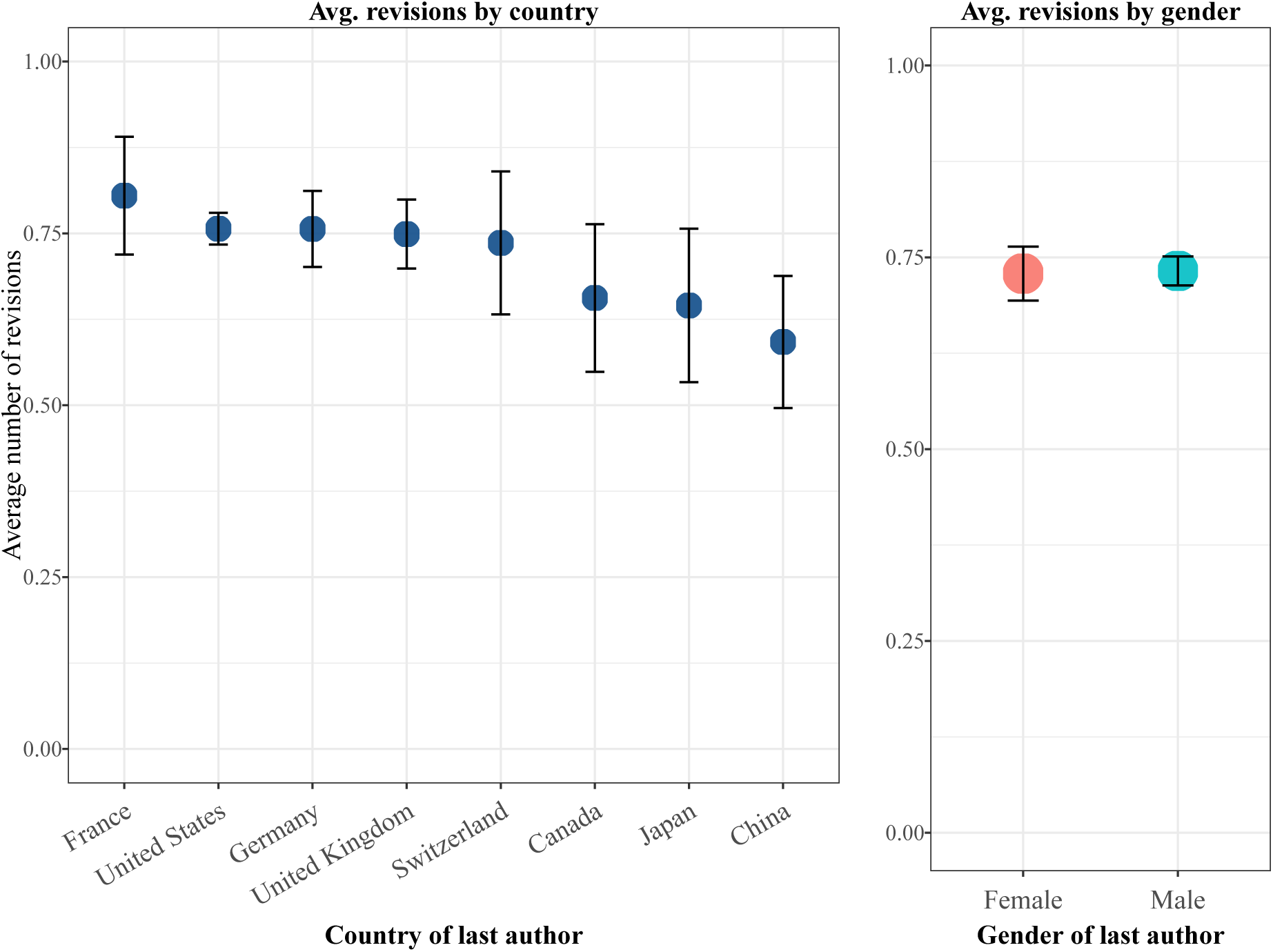
Number of revisions by author gender and country of affiliation. Average number of revisions a full submissions undergoes before a final decision of accept or reject is made. In this case, zero revisions occurs when a full submission is accepted or rejected without a request for any revisions. The dataset records at maximum two revisions, though only a small number of manuscripts remain in revision after two submissions (see Fig 1). For this figure, we only include manuscripts for which a final decision is made after zero, one, or two revisions. The left panel shows differences in the average number of revisions by the country of the last author. The right shows the average revisions by the gender of the last author. Code to reproduce this figure can be found on the linked Github repository at the path figures/revision_information/average_revisions.rmd.

**S2 Fig.**
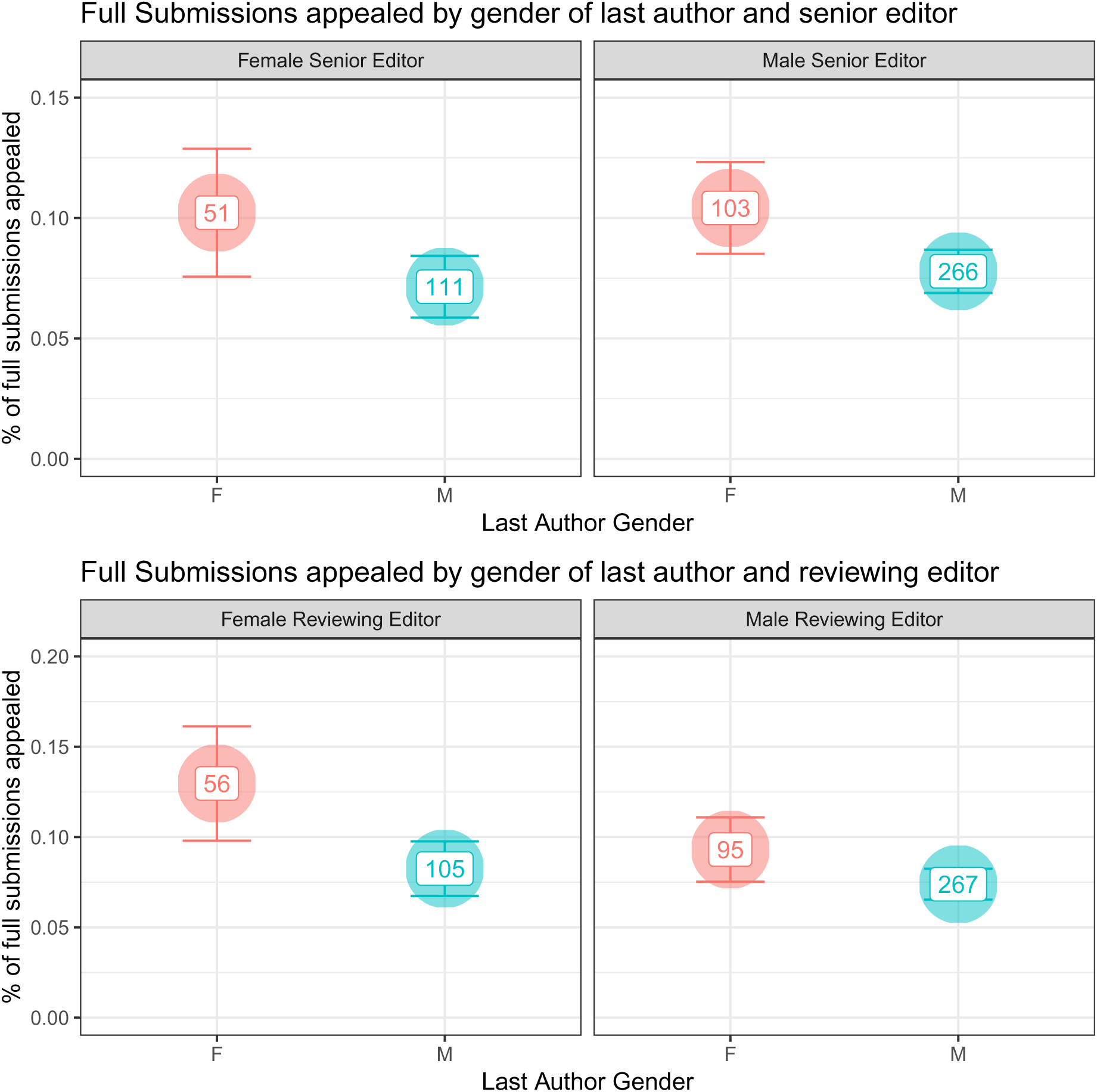
Number of appeals by gender of author and reviewing editor. Count of submissions appealed, at any review stage, by the gender of the last author gender and Senior Editor (top) and reviewing editor (bottom). Code to reproduce this figure can be found on the linked Github repository at the path figures/appeals/gender_and_appeals.rmd.

**S3 Fig.**
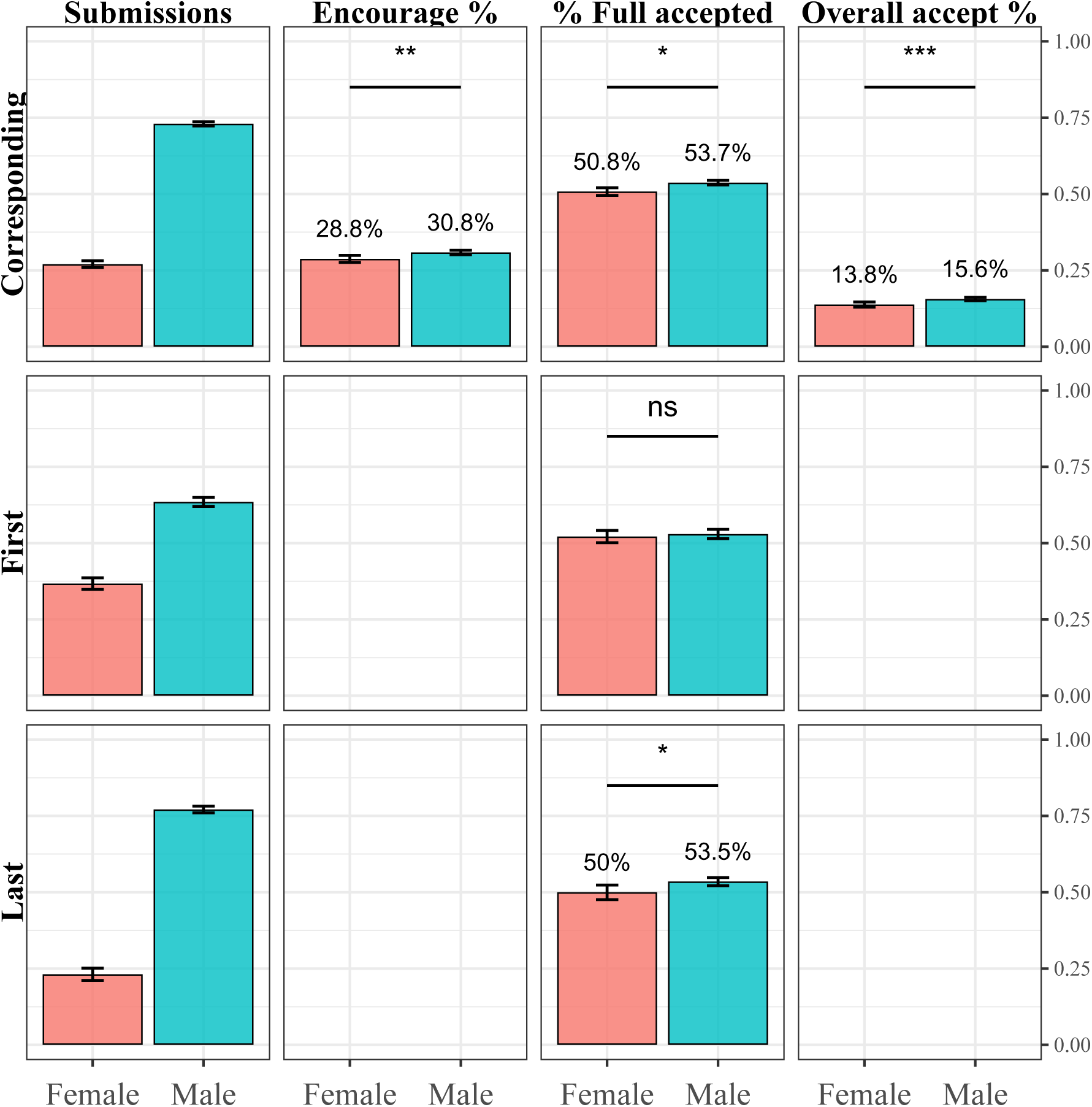
Submission and success rates by gender of corresponding, first, and last author. Proportion of initial submissions, encourage rate, overall acceptance rate, and acceptance rate of full submissions by the gender of the corresponding author, first author, and last author. Gender data is unavailable for first and last authors of initial submissions that were never submitted as full submissions, therefore these cells remain blank. Authors whose gender is unknown are excluded from analysis. Vertical error bars indicate 95% confidence intervals of the proportion of submitted, encouraged, and accepted initial and full submissions. Asterisks indicate significance level of *χ*^2^ tests of independence of frequency of encourage and acceptance by gender; “***” = *p <* 0.001; “**” = *p <* 0.01; “*” = *p <* 0.05; “-” = *p <* 0.1; “ns” = *p ≥* 0.1. Code to reproduce this figure can be found on the linked Github repository at the path figures/author_outcomes/sup_submission_outcomes.rmd.

**S4 Fig.**
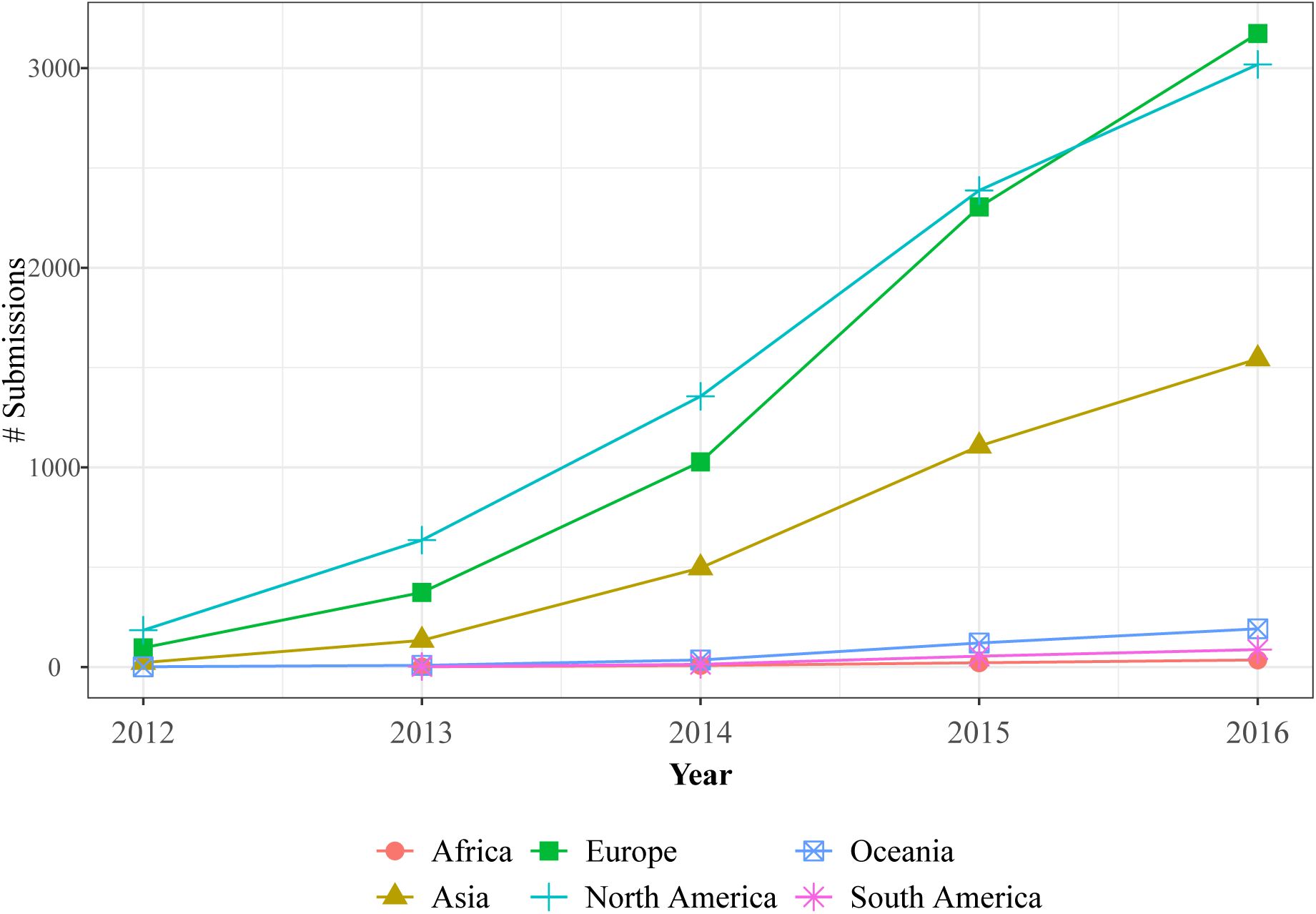
Geographic composition over time. Count of initial submissions by country of corresponding authors over time. Code to reproduce this figure can be found on the linked Github repository at the path figures/selectivity_over_time/country_composition_shift.rmd.

**S5 Fig.**
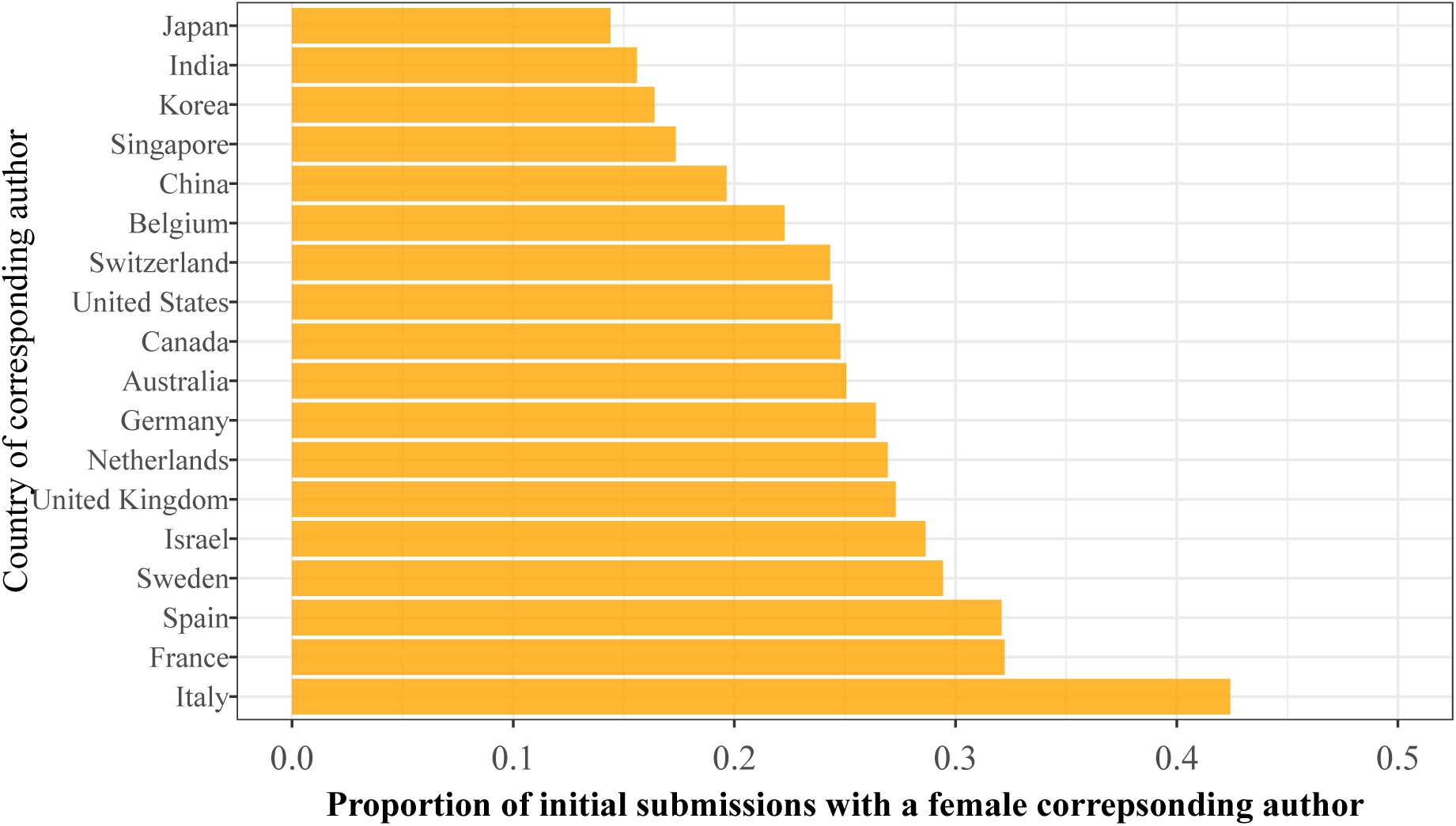
Proportion of women corresponding authors by country. Proportion of female corresponding authors on initial submissions for each country having more than 200 initial submissions during the period of study. Code to reproduce this figure can be found on the linked Github repository at the path figures/general_infromation/sup_gender_prop_by_country.rmd.

**S6 Fig.**
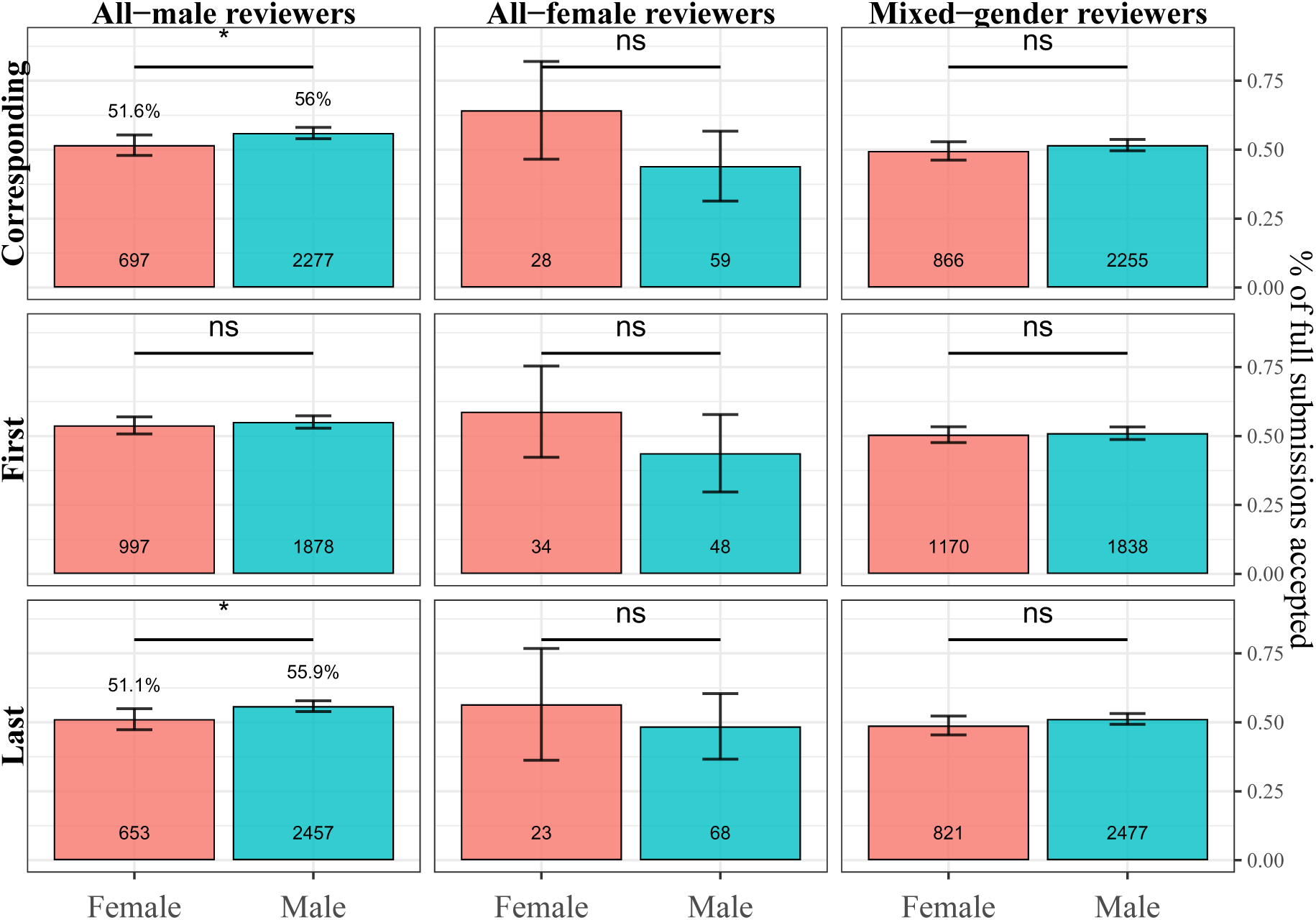
Submission and success rates by authorship role and gatekeeper gender composition. Percentage of full submissions that were accepted, shown by the gender of the corresponding, first, and last author, and by the gender composition of the peer reviewers. Text at the base of each bar indicate the number full submissions within each category of reviewer team and authorship gender. Vertical error bars indicate 95% percentile confidence intervals of the proportion of accepted full submissions. Asterisks indicate significance level of *χ*^2^ tests of independence on frequency of acceptance by gender of author given each team composition.”***” = *p <* 0.001; “**” = *p <* 0.01; “*” = *p <* 0.05; “-” = *p <* 0.1; “ns” = *p ≥* 0.1. Code to reproduce this figure can be found on the linked Github repository at the path figures/gatekeeper_author_outcomes/sup_homophily_outcomes.rmd.

**S7 Fig.**
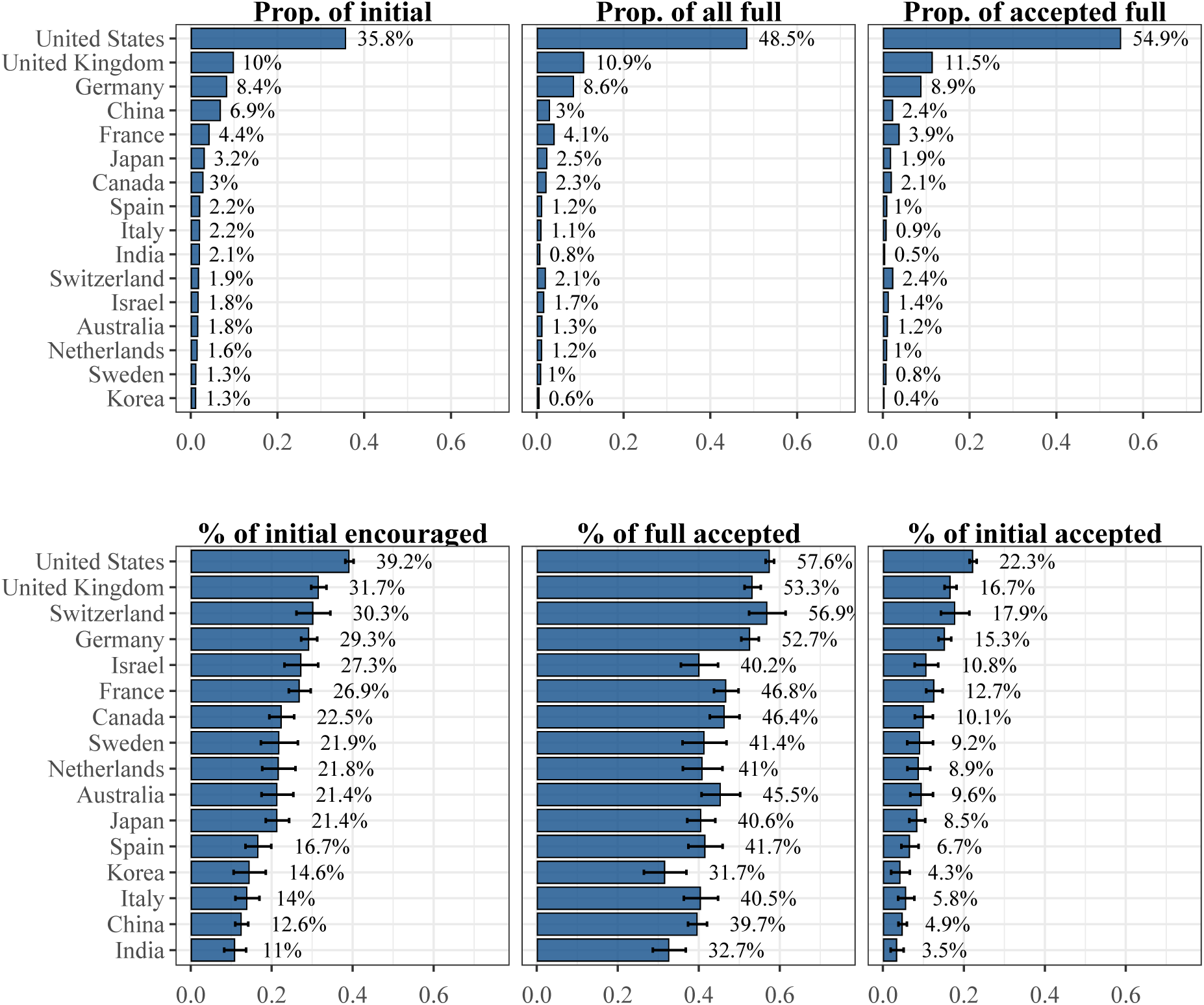
Submission and success rates by country for top 16 most prolific countries. Top: proportion of all initial submissions, encouraged initial submissions, and accepted full submissions comprised by the country of affiliation of the corresponding author for the top sixteen most prolific countries in terms of initial submissions. Bottom: acceptance rate of full submissions, encourage rate of full submissions, and overall acceptance rate of full submissions by country of affiliation of the corresponding author for the top eight more prolific countries in terms of initial submissions. Error bars on bottom panel indicate standard error of proportion of encouraged initial submissions and accepted initial and full submissions for each country. Code to reproduce this figure can be found on the linked Github repository at the path figures/author_outcomes/sup_outcomes_16_countries.rmd.

**S8 Fig.**
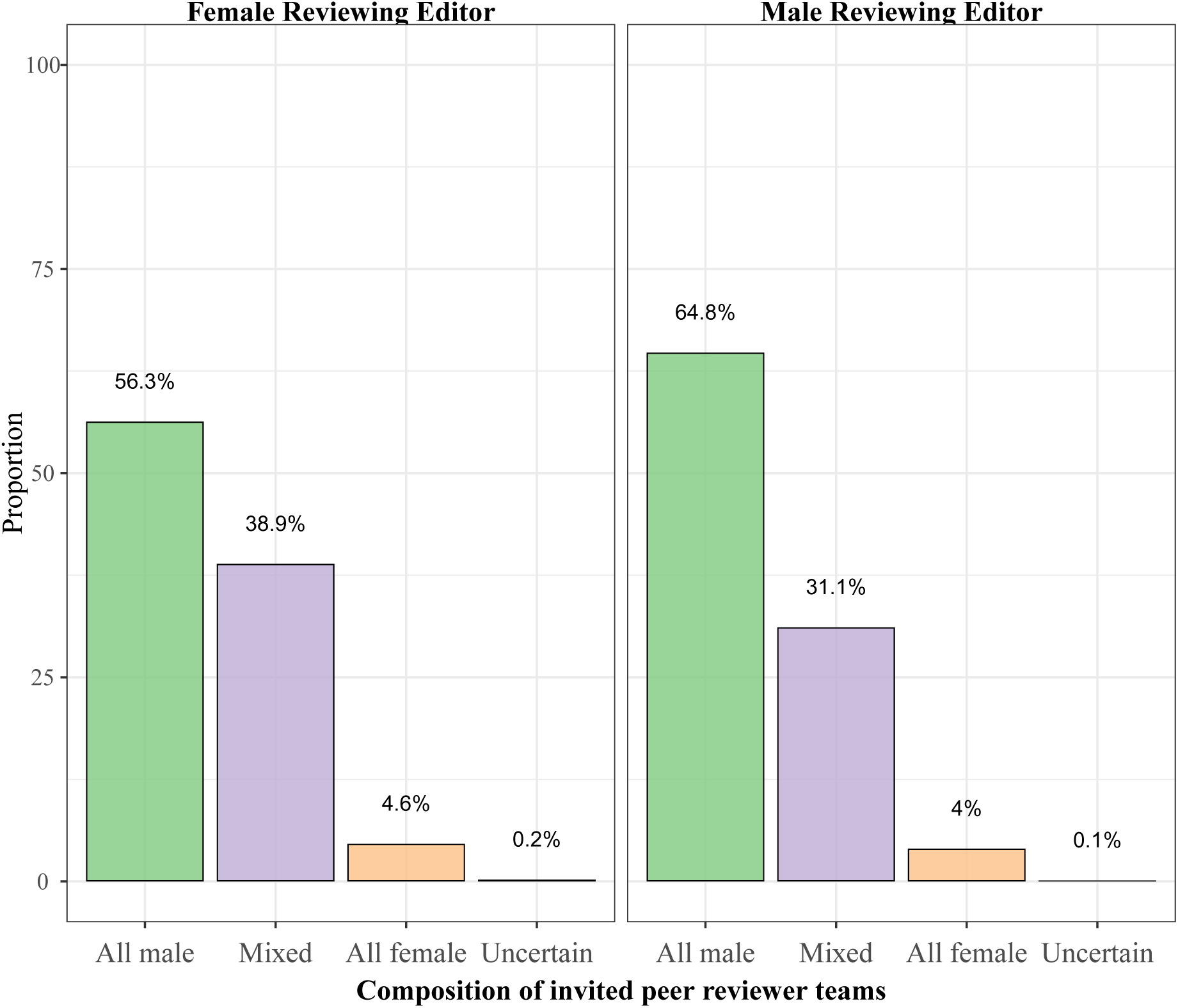
Proportion of peer reviewer team’s gender compositions by gender of the Reviewing Editor. Compositions are determined while excluding the Reviewing Editor from team membership, if they are listed as a peer reviewer. Code to reproduce this figure can be found on the linked Github repository at the path figures/gatekeeper_representation/supp_reviewing_editor_composition.rmd.

**S9 Fig.**
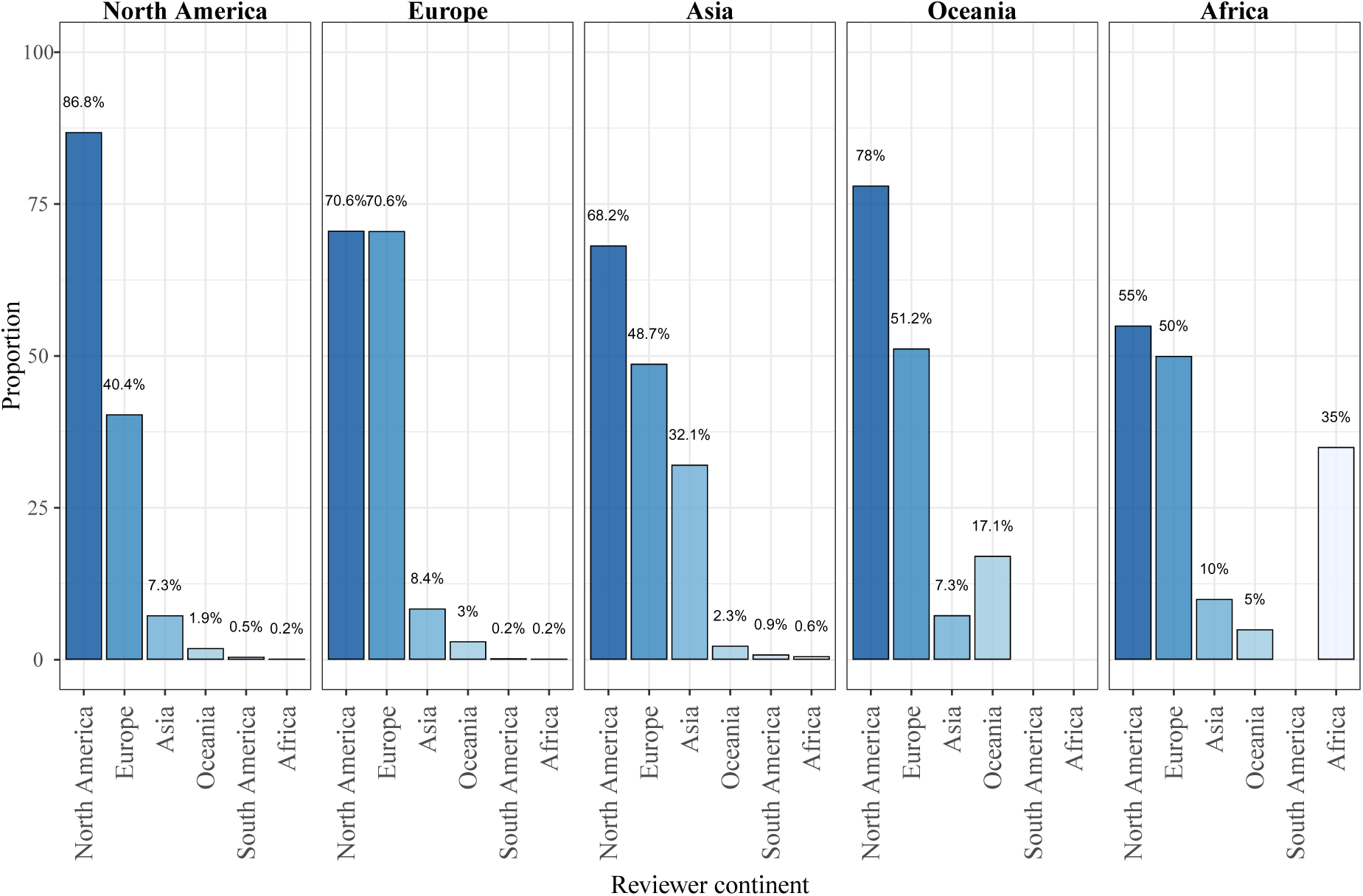
Proportion of peer review teams containing at least one peer reviewer of each continent, by continent of Reviewing Editor. Compositions are determined while excluding the Reviewing Editor from team membership, if they are listed as a peer reviewer. Code to reproduce this figure can be found on the linked Github repository at the path figures/gatekeeper_representation/reviewing_editor_continental_comp.rmd.

**S1 Table.**
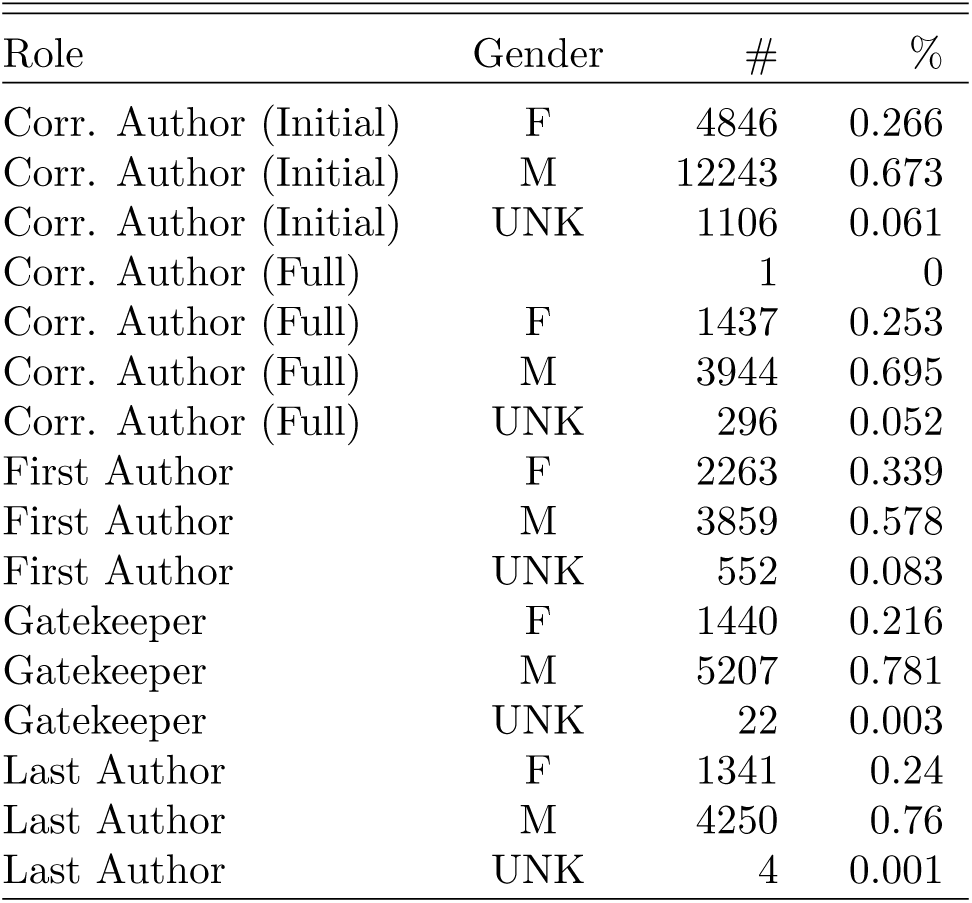
Gender demographics of *eLife*. Counts of distinct male and female corresponding authors, first authors, last authors, and gatekeepers. Includes counts on all initial and full submissions submitted between 2012 and 2017. First and last authors and gatekeepers appeared only on full submissions, whereas corresponding authors appeared on rejected or in-progress initial submissions as well. This table contains the same values as visualized in Fig 3.A.

| Role | Gender | # | % |
| --- | --- | --- | --- |
| Corr. Author (Initial) | F | 4846 | 0.266 |
| Corr. Author (Initial) | M | 12243 | 0.673 |
| Corr. Author (Initial) | UNK | 1106 | 0.061 |
| Corr. Author (Full) |  | 1 | 0 |
| Corr. Author (Full) | F | 1437 | 0.253 |
| Corr. Author (Full) | M | 3944 | 0.695 |
| Corr. Author (Full) | UNK | 296 | 0.052 |
| First Author | F | 2263 | 0.339 |
| First Author | M | 3859 | 0.578 |
| First Author | UNK | 552 | 0.083 |
| Gatekeeper | F | 1440 | 0.216 |
| Gatekeeper | M | 5207 | 0.781 |
| Gatekeeper | UNK | 22 | 0.003 |
| Last Author | F | 1341 | 0.24 |
| Last Author | M | 4250 | 0.76 |
| Last Author | UNK | 4 | 0.001 |

**S2 Table.** Summary demographic characteristics of distinct *eLife* reviewers and editors. The count of Senior Editors includes former editors, as well as the Deputy Editors and Editor-in-Chief, who also serve as Senior Editors. The count of BREs includes former editors and guest editors. Reviewers are only relevant for publications that were submitted for full review, thus leading to lower total counts. Includes all individuals involved in processing manuscripts at *eLife* between 2012 and 2017.

| <u>Reviewership</u> | <u>Female</u> |  | <u>Male</u> |  | <u>Unassigned</u> |  | <u>All</u> |
| --- | --- | --- | --- | --- | --- | --- | --- |
|  | N | % | N | % | N | % |  |
| Senior Editors | 15 | 26.3 | 42 | 73.7 | 0 | 0.0 | 57 |
| Reviewing Editors | 209 | 24.0 | 661 | 76.0 | 0.0 | 0.0 | 870 |
| Peer Reviewers | 1,526 | 21.5 | 5,572 | 78.4 | 7 | 0.1 | 7,222 |

**S3 Table.** Summary country of affiliation demographics of unique *eLife* reviewers and editors. The count of Senior Editors includes former editors, as well as the Deputy Editors and Editor-in-Chief, who also serve as Senior Editors. The count of reviewing editors includes former editors and guest editors. Reviewers are only relevant for publications that were submitted for full review, thus leading to lower total counts than the number of initial submissions. Includes all individuals involved in processing manuscripts at *eLife* between 2012 and 2017.

| Country | # Peer Rev. | % Peer Rev. | # Rev. Editor | % Rev. Editor | # Sen. Editor | % Sen. Editor |
| --- | --- | --- | --- | --- | --- | --- |
| United States | 11,313 | 0.600 | 536 | 0.620 | 32 | 0.561 |
| United Kingdom | 1,896 | 0.101 | 88 | 0.102 | 7 | 0.123 |
| Germany | 1,416 | 0.075 | 69 | 0.080 | 6 | 0.105 |
| Canada | 627 | 0.033 | 22 | 0.025 | 3 | 0.053 |
| Switzerland | 444 | 0.024 | 19 | 0.022 | 2 | 0.035 |
| China | 140 | 0.007 | 10 | 0.012 | 2 | 0.035 |
| Israel | 214 | 0.011 | 19 | 0.022 | 1 | 0.018 |
| Netherlands | 270 | 0.014 | 11 | 0.013 | 1 | 0.018 |
| Spain | 201 | 0.011 | 10 | 0.012 | 1 | 0.018 |
| Japan | 296 | 0.016 | 9 | 0.010 | 1 | 0.018 |
| India | 89 | 0.005 | 6 | 0.007 | 1 | 0.018 |
| France | 571 | 0.030 | 21 | 0.024 |  |  |
| Australia | 198 | 0.011 | 7 | 0.008 |  |  |
| South Africa | 28 | 0.001 | 5 | 0.006 |  |  |
| Austria | 118 | 0.006 | 4 | 0.005 |  |  |
| Belgium | 114 | 0.006 | 3 | 0.003 |  |  |
| Finland | 82 | 0.004 | 3 | 0.003 |  |  |
| Italy | 133 | 0.007 | 3 | 0.003 |  |  |
| Singapore | 82 | 0.004 | 3 | 0.003 |  |  |
| Thailand | 16 | 0.001 | 3 | 0.003 |  |  |
| Denmark | 78 | 0.004 | 2 | 0.002 |  |  |
| Korea | 59 | 0.003 | 2 | 0.002 |  |  |
| Estonia | 2 | 0.0001 | 1 | 0.001 |  |  |
| Hong Kong | 7 | 0.0004 | 1 | 0.001 |  |  |
| Hungary | 20 | 0.001 | 1 | 0.001 |  |  |
| Ireland | 38 | 0.002 | 1 | 0.001 |  |  |
| Kenya | 7 | 0.0004 | 1 | 0.001 |  |  |
| Mexico | 23 | 0.001 | 1 | 0.001 |  |  |
| New Zealand | 19 | 0.001 | 1 | 0.001 |  |  |
| Poland | 26 | 0.001 | 1 | 0.001 |  |  |
| Sweden | 128 | 0.007 | 1 | 0.001 |  |  |
| Albania | 2 | 0.0001 |  |  |  |  |
| Andorra | 2 | 0.0001 |  |  |  |  |
| Argentina | 21 | 0.001 |  |  |  |  |
| Brazil | 9 | 0.0005 |  |  |  |  |
| Chile | 10 | 0.001 |  |  |  |  |
| Croatia | 3 | 0.0002 |  |  |  |  |
| Czech Rep. | 8 | 0.0004 |  |  |  |  |
| Greece | 15 | 0.001 |  |  |  |  |
| Guyana | 2 | 0.0001 |  |  |  |  |
| Iceland | 2 | 0.0001 |  |  |  |  |
| Madagascar | 2 | 0.0001 |  |  |  |  |
| Malaysia | 2 | 0.0001 |  |  |  |  |
| Monaco | 1 | 0.0001 |  |  |  |  |
| Norway | 20 | 0.001 |  |  |  |  |
| Portugal | 55 | 0.003 |  |  |  |  |
| Puerto Rico | 2 | 0.0001 |  |  |  |  |
| Russia | 1 | 0.0001 |  |  |  |  |
| Saudi Arabia | 2 | 0.0001 |  |  |  |  |
| Slovenia | 1 | 0.0001 |  |  |  |  |
| Taiwan | 18 | 0.001 |  |  |  |  |
| Turkey | 4 | 0.0002 |  |  |  |  |
| United Arab Emirates | 3 | 0.0002 |  |  |  |  |
| Uruguay | 2 | 0.0001 |  |  |  |  |
| Vietnam | 1 | 0.0001 |  |  |  |  |

**S4 Table.** Geographic demographics of *eLife*. Counts of distinct corresponding authors, first authors, last authors, and gatekeepers, by continent of affiliation. Includes counts on all initial and full submissions submitted between 2012 and 2017. First and last authors and gatekeepers appeared only on full submissions, whereas corresponding authors appeared on rejected or in-progress initial submissions as well. This table contains the same values as visualized in Fig 3.B.

| Role | Continent | # | % |
| --- | --- | --- | --- |
| Corr. Author (Initial) | Africa | 61 | 0.003 |
| Corr. Author (Initial) | Asia | 3238 | 0.178 |
| Corr. Author (Initial) | Europe | 7264 | 0.399 |
| Corr. Author (Initial) | North America | 7045 | 0.387 |
| Corr. Author (Initial) | Oceania | 399 | 0.022 |
| Corr. Author (Initial) | South America | 188 | 0.01 |
| Corr. Author (Full) | Africa | 10 | 0.002 |
| Corr. Author (Full) | Asia | 624 | 0.11 |
| Corr. Author (Full) | Europe | 2078 | 0.366 |
| Corr. Author (Full) | North America | 2854 | 0.503 |
| Corr. Author (Full) | Oceania | 95 | 0.017 |
| Corr. Author (Full) | South America | 17 | 0.003 |
| First Author | Africa | 14 | 0.002 |
| First Author | Asia | 751 | 0.113 |
| First Author | Europe | 2373 | 0.356 |
| First Author | North America | 3412 | 0.511 |
| First Author | Oceania | 102 | 0.015 |
| First Author | South America | 22 | 0.003 |
| Gatekeeper | Africa | 17 | 0.003 |
| Gatekeeper | Asia | 378 | 0.057 |
| Gatekeeper | Europe | 2162 | 0.324 |
| Gatekeeper | North America | 3992 | 0.599 |
| Gatekeeper | Oceania | 98 | 0.015 |
| Gatekeeper | South America | 22 | 0.003 |
| Last Author | Africa | 13 | 0.002 |
| Last Author | Asia | 619 | 0.111 |
| Last Author | Europe | 2063 | 0.369 |
| Last Author | North America | 2789 | 0.498 |
| Last Author | Oceania | 94 | 0.017 |
| Last Author | South America | 17 | 0.003 |

**S5 Table.** Submissions and proportion of author/gatekeeper homogeny by country. Includes number of full submissions submitted with corresponding authors from each of 20 countries, and proportion of these full submissions with the condition of author/reviewer homogeny such that at least one involved gatekeeper from the same country. Countries listed are in order of the proportion of author/reviewer homogeny, and contain the top 20 countries with the highest homogeny.

| Country | # Submissions | # Homogeneity | % Country Homogeneity |
| --- | --- | --- | --- |
| United States | 3605 | 3185 | 0.883 |
| United Kingdom | 803 | 236 | 0.294 |
| Germany | 641 | 168 | 0.262 |
| Mexico | 5 | 1 | 0.2 |
| Korea | 45 | 8 | 0.178 |
| Canada | 176 | 27 | 0.153 |
| Japan | 184 | 19 | 0.103 |
| Australia | 101 | 10 | 0.099 |
| China | 233 | 23 | 0.099 |
| Switzerland | 163 | 16 | 0.098 |
| Ireland | 11 | 1 | 0.091 |
| South Africa | 11 | 1 | 0.091 |
| France | 310 | 28 | 0.09 |
| Poland | 12 | 1 | 0.083 |
| Belgium | 41 | 3 | 0.073 |
| Finland | 14 | 1 | 0.071 |
| Norway | 14 | 1 | 0.071 |
| India | 59 | 4 | 0.068 |
| Denmark | 32 | 2 | 0.062 |

**S6 Table.** Model coefficients of initial submissions—author characteristics: Odds ratio, associated confidence intervals, and model diagnostics for logistic regression model using the encouragement of initial submission as a response variable. Predictor variables include control variables of the submission year and type, and variables capturing author characteristics. For continent of affiliation, “North America” was used as the reference level. For submission type, “RA” (research article) was used as the reference level; the submission type “SR” means “Short Reports”, and “TR” means “Tools and Resources”. This table contains the same values as visualized in Fig 5.A.

|  | ENCOURAGED |
| --- | --- |
|  | *logistic* |
| Submission Year | .918***<br>(.894,.942) |
| Submission Type = SR | .742***<br>(.638,.847) |
| Submission Type = TR | .740***<br>(.567,.913) |
| Corr. Author is Male | 1.118**<br>(1.051,1.185) |
| Corr. Author Gender UNK | .932<br>(.795,1.070) |
| Corr. Author Inst. Top | 1.726***<br>(1.663,1.789) |
| Corr. Author from Africa | .535*<br>(-.018,1.088) |
| Corr. Author from Asia | .395***<br>(.301,.488) |
| Corr. Author from Europe | .676***<br>(.611,.740) |
| Corr. Author from Oceania | .559***<br>(.336,.783) |
| Corr. Author from South America | .205***<br>(-.269,.679) |
| Constant | .638***<br>(.526,.749) |
| Observations | 23,615 |
| Log Likelihood | -13,778.170 |
| Akaike Inf. Crit. | 27,580.330 |
| Notes: | *P < .05<br>**P < .01<br>***P < .001 |

**S7 Table.** Model coefficients of full submissions—author characteristics: Odds ratio, associated confidence intervals, and model diagnostics for logistic regression model using the acceptance of full submission as a response variable. Predictor variables include control variables of the submission year and type, and variables capturing author characteristics. For continent of affiliation, “North America” was used as the reference level. For submission type, “RA” (research article) was used as the reference level; the submission type “SR” means “Short Reports”, and “TR” means “Tools and Resources”. This table contains the same values as visualized in Fig 5.B.

|  | ACCEPTED<br>*logistic* |
| --- | --- |
| Submission Year | .888***<br>(.847,.929) |
| Submission Type = SR | .897<br>(.711,1.082) |
| Submission Type = TR | 1.117<br>(.800,1.434) |
| First Author is Male | 1.022<br>(.914,1.129) |
| First Author is Unknown Gender | 1.033<br>(.840,1.226) |
| Last Author is Male | 1.145*<br>(1.027,1.263) |
| Last Author Inst. Top | 1.379***<br>(1.272,1.486) |
| Last Author from Africa | 1.484<br>(.464,2.503) |
| Last Author from Asia | .585***<br>(.408,.763) |
| Last Author from Europe | .860**<br>(.749,.972) |
| Last Author from Oceania | .906<br>(.490,1.323) |
| Last Author from South America | .839<br>(-.098,1.776) |
| Constant | 1.430***<br>(1.230,1.629) |
| Observations | 6,461 |
| Log Likelihood | -4,390.813 |
| Akaike Inf. Crit. | 8,807.626 |
| Notes: | *P < .05<br>**P < .01<br>***P < .001 |

**S8 Table.** Model coefficients of initial submissions—author characteristics and homogeny: Odds ratio, associated confidence intervals, and model diagnostics for logistic regression model using the encouragement of initial submission as a response variable. Predictor variables include control variables of the submission year and type, and variables capturing author characteristics and author-reviewer homogeny. For continent of affiliation, “North America” was used as the reference level. For submission type, “RA” (research article) was used as the reference level; the submission type “SR” means “Short Reports”, and “TR” means “Tools and Resources”.

|  | ENCOURAGED |
| --- | --- |
|  | *logistic* |
| Submission Year | .918***<br>(.894,.942) |
| Submission Type = SR | .742***<br>(.638,.847) |
| Submission Type = TR | .741***<br>(.568,.914) |
| Corr. Author is Male | 1.115**<br>(1.048,1.182) |
| Corr. Author is Unknown Gender | .930<br>(.792,1.068) |
| Corr. Author Inst. Top | 1.709***<br>(1.645,1.772) |
| Corr. Author from Africa | .579<br>(.021,1.137) |
| Corr. Author from Asia | .443***<br>(.337,.549) |
| Corr. Author from Europe | .800***<br>(.724,.877) |
| Corr. Author from Oceania | .570***<br>(.328,.813) |
| Corr. Author from South America | .225***<br>(-.254,.703) |
| Corr. Author-Editor Geo. Distance | 1.022***<br>(1.010,1.034) |
| Corr. Author-Editor Geo. Distance = 0 | 1.560***<br>(1.448,1.673) |
| Constant | .465***<br>(.320,.610) |
| Observations | 23,615 |
| Log Likelihood | -13,742.830 |
| Akaike Inf. Crit. | 27,513.650 |
| Notes: | *P < .05<br>**P < .01<br>***P < .001 |

**S9 Table.** Model coefficients of regressions on full submissions: Odds ratio, associated confidence intervals, and model diagnostics for logistic regression model using the acceptance of full submission as the response variable. Control variables include the submission year, submission type, last author institutional prestige, and the gender of the first author. Other predictor variables include the gender of the last author, continent of affiliation of the last author, gender-composition of the reviewers, the last author-reviewers geographic distance, and variables attempting to capture the gender equity by reviewer-team composition group. Five models are presented: the first (Main Effects) shows only the main effects for the model including all full submissions without any additional manipulation or variables (1); the second model (2, Standard Interaction) models the main effects as well as an interaction term between last author gender and the gender composition of the reviewer team (an ANOVA table for this model has been provided in S11 Table; the next two models were separately trained on only submissions reviewed by all-male reviewer teams (3) and only submission trained on mixed-gender reviewer teams (4), respectively; the last model (5) models gender equity between reviewer-composition groups using a new variable with all combinations of author and reviewer gender (see Fig 7). Columns (1) and (5) contain the same values as Fig 7A and Fig 7.B, respectively. For continent of affiliation, “North America” was used as the reference level. For submission type, “RA” (research article) was used as the reference level; the submission type “SR” means “Short Reports”, and “TR” means “Tools and Resources”. For the combination variable of last author gender and reviewer team composition, we held “last author female, all rev. male” as the reference level. Missing cells indicates that the corresponding variable was not part of that model.

|  | ACCEPTED |  |
| --- | --- | --- |
|  | *logistic* |  |
|  | All Male<br>1 | Mixed-Gender<br>2 |
| Submission Year | .907**<br>(.848,.966) | .881***<br>(.823,.940) |
| Submission Type = SR | .993<br>(.727,1.259) | .770<br>(.503,1.038) |
| Submission Type = TR | 1.035<br>(.574,1.496) | 1.139<br>(.692,1.586) |
| First Author is Male | 1.034<br>(.875,1.193) | 1.022<br>(.873,1.172) |
| First Author is Unknown Gender | 1.163<br>(.869,1.456) | .967<br>(.704,1.230) |
| Last Author Inst. Top | 1.519***<br>(1.362,1.676) | 1.330***<br>(1.180,1.480) |
| Last Author Male | 1.228*<br>(1.051,1.405) | 1.088<br>(.926,1.249) |
| Last author from Africa | 2.212<br>(.477,3.948) | 2.276<br>(.972,3.581) |
| Last author from Asia | .758<br>(.447,1.068) | .851<br>(.551,1.152) |
| Last author from Europe | 1.020<br>(.835,1.205) | .951<br>(.776,1.125) |
| Last author from Oceania | .974<br>(.312,1.636) | 2.516**<br>(1.826,3.205) |
| Last author from South America | .975<br>(-.543,2.492) | 1.656<br>(.390,2.923) |
| Sum of geo. distance (1000s km) | .992<br>(.983,1.001) | .982***<br>(.973,.991) |
| Sum of geo. distance is zero | 1.240<br>(.996,1.483) | .797<br>(.558,1.037) |
| All Reviewers Male | 1.271<br>(.940,1.601) | 1.872***<br>(1.558,2.187) |
| Observations | 3,090 | 3,280 |
| Log Likelihood | -2,074.757 | -2,228.574 |
| Akaike Inf. Crit. | 4,179.513 | 4,487.148 |
| Notes: | *P < .05<br>**P < .01<br>***P < .001 |  |

**S10 Table.** Continent-level table of initial and full submission counts: The number of initial and full submissions for each of seven continents including Antarctica (which was excluded from other analysis).

|  | Continent | # Initial submissions | # Full submissions |
| --- | --- | --- | --- |
| 1 | North America | 9591 | 3785 |
| 2 | Europe | 9106 | 2527 |
| 3 | Asia | 4382 | 735 |
| 4 | Oceania | 472 | 107 |
| 5 | South America | 215 | 20 |
| 6 | Africa | 78 | 17 |
| 7 | Antarctica | 1 | NA |

**S11 Table.** ANOVA table for author-reviewer interaction model: Results of ANOVA test run on the fitted model containing main effects for author and reviewer characteristics for full submissions as well as the interaction between last author gender and reviewer team composition.

|  | term | df | Deviance | Resid..Df | Resid..Dev | p.value |
| --- | --- | --- | --- | --- | --- | --- |
| 1 | Submission Year | 1 | 47.997 | 6459 | 8889.773 | <0.0001 |
| 2 | Submission Type | 2 | 2.397 | 6457 | 8887.377 | 0.30172 |
| 3 | First Author Gender | 2 | 0.306 | 6455 | 8887.071 | 0.85814 |
| 4 | Last Author Inst. Prestige | 1 | 62.855 | 6454 | 8824.216 | <0.0001 |
| 5 | Last Author Gender | 1 | 5.194 | 6453 | 8819.022 | 0.02266 |
| 6 | Last author Continent | 5 | 37.397 | 6448 | 8781.626 | <0.0001 |
| 7 | Last Author-Reviewers Geographic Distance | 1 | 22.679 | 6447 | 8758.946 | <0.0001 |
| 8 | Sum of geo. distance is zero | 1 | 0.018 | 6446 | 8758.928 | 0.89338 |
| 9 | Reviewer Gender Composition | 2 | 7.797 | 6444 | 8751.131 | 0.02027 |
| 10 | Last Author Gender*Reviewer Gender Composition | 2 | 1.767 | 6442 | 8749.365 | 0.4134 |

**S11 Table. Model coefficients of full submissions—author characteristics and reviewing-editor only homogeny:** Odds ratio, associated confidence intervals, and model diagnostics for logistic regression model using the encouragement of full submission as a response variable. Predictor variables include control variables of the submission year and type, and variables capturing author characteristics and homogeny between the author and reviewing editor only. For continent of affiliation, “North America” was used as the reference level. For submission type, “RA” (research article) was used as the reference level; the submission type “SR” means “Short Reports”, and “TR” means “Tools and Resources”. This regression models gender equity between reviewer composition groups using a new variable containing all combinations of last author gender and reviewer team composition; for this new categorical variable, we used “last author female - female rev. editor” as the reference level.

**Table 1.**

|  | ACCEPTED |
| --- | --- |
| Submission Year | .897***<br>(.856,.939) |
| Submission Type = SR | .890<br>(.703,1.078) |
| Submission Type = TR | 1.090<br>(.767,1.413) |
| First Author is Male | 1.010<br>(.901,1.119) |
| First Author is Unknown Gender | 1.057<br>(.862,1.253) |
| Last Author Inst. Top 50 | 1.383***<br>(1.275,1.492) |
| Last author from Africa | 2.239<br>(1.201,3.278) |
| Last author from Asia | .805<br>(.586,1.024) |
| Last author from Europe | 1.002<br>(.863,1.141) |
| Last author from Oceania | 1.520<br>(1.039,2.000) |
| Last author from South America | 1.194<br>(.246,2.142) |
| Dist. between author and rev. editor (1000km) | 1.018<br>(.990,1.045) |
| Sum of author-reviewer distance (1000km) | .984***<br>(.975,.993) |
| Total dist. between author and reviewers is zero | .977<br>(.792,1.162) |
| Dist. between author and rev. editor is zero | 1.095<br>(.882,1.308) |
| Last author female - male rev. editor | 1.204<br>(.975,1.433) |
| Last author male - female rev. editor | 1.178<br>(.955,1.400) |
| Last author male - male rev. editor | 1.352**<br>(1.148,1.556) |
| All Female Reviewers | .962<br>(.706,1.218) |
| Mixed Reviewers | .951<br>(.843,1.060) |
| Constant | 1.306<br>(.991,1.621) |
| Observations | 6,320 |
| Log Likelihood | -4,280.736 |
| Akaike Inf. Crit. | 8,603.471 |
| Notes: | *P < .05<br>**P < .01<br>***P < .001 |

## Acknowledgments

We are grateful for the editing and feedback provided by Susanna Richmond (Senior Manager at *eLife*), Mark Patterson (Executive Director at *eLife*), Eve Marder, Anna Akhmanova, and Detlef Weigel (Deputy Editors at *eLife*). We are also grateful for the work of James Gilbert (Production Editor at *eLife*) for extracting the data used in this analysis. This work was partially supported by a grant from the National Science Foundation (SciSIP #1561299).

## Ethics statement

This research underwent expedited review by the Institutional Review Board at Indiana University Bloomington and was determined to be exempt (Protocol #: 1707327848).

## References

1. Merton RK. The Matthew Effect in Science: The reward and communication systems of science are considered. Science. 1968;159(3810):56–63. doi:10.1126/science.159.3810.56.

2. Walker R, Barros B, Conejo R, Neumann K, Telefont M. Personal attributes of authors and reviewers, social bias and the outcomes of peer review: a case study. F1000Research. 2015;doi:10.12688/f1000research.6012.2.

3. Lee CJ, Sugimoto CR, Zhang G, Cronin B. Bias in peer review. Journal of the American Society for Information Science and Technology. 2013;64(1):2–17. doi:10.1002/asi.22784.

4. Pinholster G. Journals and funders confront implicit bias in peer review. Science. 2016;352(6289):1067–1068. doi:10.1126/science.352.6289.1067.

5. Kaatz A, Gutierrez B, Carnes M. Threats to objectivity in peer review: the case of gender. Trends in pharmacological sciences. 2014;35(8):371–373. doi:10.1016/j.tips.2014.06.005.

6. Lariviè re V, Ni C, Gingras Y, Cronin B, Sugimoto CR. Bibliometrics: Global gender disparities in science. Nature News. 2013;504(7479):211. doi:10.1038/504211a.

7. Women, Minorities, and Persons with Disabilities in Science and Engineering: 2017. Arlington, VA: National Science Foundation, National Center for Science and Engineering Statistics.; 2017.

8. Gender in the Global Research Landscape. Elsevier; 2017. Available from: https://www.elsevier.com/data/assets/pdf_file/0008/265661/ElsevierGenderReport_final_for-web.pdf.

9. West JD, Jacquet J, King MM, Correll SJ, Bergstrom CT. The Role of Gender in Scholarly Authorship. PLOS ONE. 2013;8(7):e66212. doi:10.1371/journal.pone.0066212.

10. Larivière V, Sugimoto CR. The end of gender disparities in science? If only it were true…; 2017. Available from: https://www.cwts.nl:443/blog?article=n-q2z294&title=the-end-of-gender-disparities-in-science-if-only-it-were-true.

11. Bendels MHK, Müller R, Brueggmann D, Groneberg DA. Gender disparities in high-quality research revealed by Nature Index journals. PLOS ONE. 2018;13(1):e0189136. doi:10.1371/journal.pone.0189136.

12. Bernard C. Editorial: Gender Bias in Publishing: Double-Blind Reviewing as a Solution? eNeuro. 2018;5(3):ENEURO.0225–18.2018. doi:10.1523/ENEURO.0225-18.2018.

13. Shen YA, Webster JM, Shoda Y, Fine I. Persistent Underrepresentation of Women’s Science in High Profile Journals. bioRxiv. 2018; p. 275362. doi:10.1101/275362.

14. Nature’s under-representation of women. Nature. 2018;558(7710):344. doi:10.1038/d41586-018-05465-7.

15. King DA. The scientific impact of nations. Nature. 2004;430:311.

16. White KE, Robbins C, Khan B, Freyman C. Science and Engineering Publication Output Trends: 2014 Shows Rise of Developing Country Output while Developed Countries Dominate Highly Cited Publications. Arlington, VA: National Center for Science and Engineering Statistics; 2017. NSF 18–300.

17. Wennerås C, Wold A. Nepotism and sexism in peer-review. Nature. 1997;387:341.

18. Li D. Gender Bias in NIH Peer Review: Does it Exist and Does it Matter?; 2011. Available from: http://citeseerx.ist.psu.edu/viewdoc/download?doi=10.1.1.571.8269&rep=rep1&type=pdf.

19. Tamblyn R, Girard N, Qian CJ, Hanley J. Assessment of potential bias in research grant peer review in Canada. CMAJ. 2018;190(16):E489–E499. doi:10.1503/cmaj.170901.

20. Witteman HO, Hendricks M, Straus S, Tannenbaum C. Female grant applicants are equally successful when peer reviewers assess the science, but not when they assess the scientist. bioRxiv. 2017; p. 232868. doi:10.1101/232868.

21. Grant J, Burden S, Breen G. No evidence of sexism in peer review. Nature. 1997;390(6659):438. doi:10.1038/37213.

22. Gilbert JR, Williams ES, Lundberg GD. Is there gender bias in JAMA’s peer review process? JAMA. 1994;272(2):139–142.

23. Mutz R, Bornmann L, Daniel HD. Does Gender Matter in Grant Peer Review?: An Empirical Investigation Using the Example of the Austrian Science Fund. Zeitschrift Fur Psychologie. 2012;220(2):121–129. doi:10.1027/2151-2604/a000103.

24. Beck R, Halloin V. Gender and research funding success: Case of the Belgian F.R.S.-FNRS. Research Evaluation. 2017;26(2):115–123. doi:10.1093/reseval/rvx008.

25. Edwards HA, Schroeder J, Dugdale HL. Gender differences in authorships are not associated with publication bias in an evolutionary journal. PLOS ONE. 2018;13(8):e0201725. doi:10.1371/journal.pone.0201725.

26. Coates R, Sturgeon B, Bohannan J, Pasini E. Language and publication in Cardiovascular Research articles. Cardiovascular Research. 2002;53(2):279–285. doi:10.1016/S0008-6363(01)00530-2.

27. Primack RB, Marrs R. Bias in the review process. Biological Conservation. 2008;141(12):2919–2920. doi:10.1016/j.biocon.2008.09.016.

28. Harris M, Marti J, Watt H, Bhatti Y, Macinko J, Darzi AW. Explicit Bias Toward High-Income-Country Research: A Randomized, Blinded, Crossover Experiment Of English Clinicians. Health Affairs (Project Hope). 2017;36(11):1997–2004. doi:10.1377/hlthaff.2017.0773.

29. Primack RB, Ellwood E, Miller-Rushing AJ, Marrs R, Mulligan A. Do gender, nationality, or academic age affect review decisions? An analysis of submissions to the journal Biological Conservation. Biological Conservation. 2009;142(11):2415–2418. doi:10.1016/j.biocon.2009.06.021.

30. Berger J, Fisek HF, Normal RZ, Zelditch N. Status characteristics and social interaction. New York: Elsevier; 1977.

31. Correll SJ, Ridgeway CL. Expectation States Theory. In: Handbook of Social Psychology. Handbooks of Sociology and Social Research. Springer, Boston, MA; 2006. p. 29–51. Available from: https://link.springer.com/chapter/10.1007/0-387-36921-X_2.

32. Podolny JM. Status Signals: A Sociological Study of Market Competition. Princeton, N.J Woodstock: Princeton University Press; 2008.

33. Long JS, Fox MF. Scientific Careers: Universalism and Particularism. Annual Review of Sociology. 1995;21:45–71.

34. Pfeffer J, Leong A, Strehl K. Paradigm Development and Particularism: Journal Publication in Three Scientific Disciplines. Social Forces. 1977;55(4):938–951. doi:10.2307/2577563.

35. Cole S. The Hierarchy of the Sciences? American Journal of Sociology. 1983;89(1):111–139. doi:10.1086/227835.

36. Clauset A, Arbesman S, Larremore DB. Systematic inequality and hierarchy in faculty hiring networks. Science Advances. 2015;1(1):e1400005. doi:10.1126/sciadv.1400005.

37. Jacobs JA. Gender and the Stratification of Colleges. The Journal of Higher Education. 1999;70(2):161–187. doi:10.2307/2649126.

38. Weeden KA, Thebaud S, Gelbgiser D. Degrees of Difference: Gender Segregation of U.S. Doctorates by Field and Program Prestige. Sociological Science. 2017;4:123–150. doi:10.15195/v4.a6.

39. Sheltzer JM, Smith JC. Elite male faculty in the life sciences employ fewer women. Proceedings of the National Academy of Sciences. 2014;111(28):10107–10112. doi:10.1073/pnas.1403334111.

40. Travis GDL, Collins HM. New Light on Old Boys: Cognitive and Institutional Particularism in the Peer Review System. Science, Technology, & Human Values. 1991;16(3):322–341.

41. Teplitskiy M, Acuna D, Elamrani-Raoult A, Körding K, Evans J. The sociology of scientific validity: How professional networks shape judgement in peer review. Research Policy. 2018;doi:10.1016/j.respol.2018.06.014.

42. Demarest B, Freeman G, Sugimoto CR. The reviewer in the mirror: examining gendered and ethnicized notions of reciprocity in peer review. Scientometrics. 2014;101(1):717–735. doi:10.1007/s11192-014-1354-z.

43. Link AM. US and non-US submissions: an analysis of reviewer bias. JAMA. 1998;280(3):246–247.

44. Helmer M, Schottdorf M, Neef A, Battaglia D. Research: Gender bias in scholarly peer review. eLife. 2017;6:e21718. doi:10.7554/eLife.21718.

45. Bagues M, Sylos-Labini M, Zinovyeva N. Does the Gender Composition of Scientific Committees Matter? American Economic Review. 2017;107(4):1207–1238. doi:10.1257/aer.20151211.

46. Ceci SJ, Williams WM. Understanding current causes of women’s underrepresentation in science. Proceedings of the National Academy of Sciences. 2011;108(8):3157–3162. doi:10.1073/pnas.1014871108.

47. Ceci SJ, Ginther DK, Kahn S, Williams WM. Women in Academic Science: A Changing Landscape. Psychological Science in the Public Interest: A Journal of the American Psychological Society. 2014;15(3):75–141. doi:10.1177/1529100614541236.

48. Hengel E. Publishing while Female. Are women held to higher standards? Evidence from peer review. Faculty of Economics, University of Cambridge; 2017. 1753. Available from: https://ideas.repec.org/p/cam/camdae/1753.html.

49. Niederle M, Vesterlund L. Gender and Competition. Annual Review of Economics. 2011;3(1):601–630. doi:10.1146/annurev-economics-111809-125122.

50. May RM. The Scientific Wealth of Nations. Science. 1997;275(5301):793–796. doi:10.1126/science.275.5301.793.

51. Quan W, Chen B, Shu F. Publish or impoverish: An investigation of the monetary reward system of science in China (1999-2016). Aslib Journal of Information Management. 2017;69(5):486–502. doi:10.1108/AJIM-01-2017-0014.

52. Duszak A, Lewkowicz J. Publishing academic texts in English: A Polish perspective. Journal of English for Academic Purposes. 2008;7(2):108–120. doi:10.1016/j.jeap.2008.03.001.

53. Salager-Meyer F. Scientific publishing in developing countries: Challenges for the future. Journal of English for Academic Purposes. 2008;7(2):121–132. doi:10.1016/j.jeap.2008.03.009.

54. Yang W. Policy: Boost basic research in China. Nature News. 2016;534(7608):467. doi:10.1038/534467a.

55. Langfeldt L. The Decision-Making Constraints and Processes of Grant Peer Review, and Their Effects on the Review Outcome. Social Studies of Science. 2001;31(6):820–841. doi:10.1177/030631201031006002.

56. Lamont M. How Professors Think: Inside the Curious World of Academic Judgment. Cambridge: Harvard University Press; 2009.

57. Schekman R, Watt F, Weigel D. Scientific Publishing: The eLife approach to peer review. eLife. 2013;2:e00799. doi:10.7554/eLife.00799.

58. King SR. Peer Review: Consultative review is worth the wait. eLife. 2017;6:e32012. doi:10.7554/eLife.32012.

59. Costas R, Bordons M. Do age and professional rank influence the order of authorship in scientific publications? Some evidence from a micro-level perspective. Scientometrics. 2011;88(1):145–161. doi:10.1007/s11192-011-0368-z.

60. Macaluso B, Lariviè re V, Sugimoto T, Sugimoto CR. Is Science Built on the Shoulders of Women? A Study of Gender Differences in Contributorship. Academic Medicine: Journal of the Association of American Medical Colleges. 2016;91(8):1136–1142. doi:10.1097/ACM.0000000000001261.

61. Baerlocher MO, Newton M, Gautam T, Tomlinson G, Detsky AS. The meaning of author order in medical research. Journal of Investigative Medicine: The Official Publication of the American Federation for Clinical Research. 2007;55(4):174–180. doi:10.2310/6650.2007.06044.

62. Tscharntke T, Hochberg ME, Rand TA, Resh VH, Krauss J. Author Sequence and Credit for Contributions in Multiauthored Publications. PLOS Biology. 2007;5(1):e18. doi:10.1371/journal.pbio.0050018.

63. Winkler WE. String Comparator Metrics and Enhanced Decision Rules in the Fellegi-Sunter Model of Record Linkage; 1990. Available from: https://eric.ed.gov/?id=ED325505.

64. Harvard WorldMap;. Available from: http://worldmap.harvard.edu/.

65. Kliewer MA, DeLong DM, Freed K, Jenkins CB, Paulson EK, Provenzale JM. Peer review at the American Journal of Roentgenology: how reviewer and manuscript characteristics affected editorial decisions on 196 major papers. AJR American journal of roentgenology. 2004;183(6):1545–1550. doi:10.2214/ajr.183.6.01831545.

66. Ross JS, Gross CP, Desai MM, Hong Y, Grant AO, Daniels SR, et al. Effect of blinded peer review on abstract acceptance. JAMA. 2006;295(14):1675–1680. doi:10.1001/jama.295.14.1675.

67. Amrein K, Langmann A, Fahrleitner-Pammer A, Pieber TR, Zollner-Schwetz I. Women Underrepresented on Editorial Boards of 60 Major Medical Journals. Gender Medicine. 2011;8(6):378–387. doi:10.1016/j.genm.2011.10.007.

68. Cho AH, Johnson SA, Schuman CE, Adler JM, Gonzalez O, Graves SJ, et al. Women are underrepresented on the editorial boards of journals in environmental biology and natural resource management. PeerJ. 2014;2:e542. doi:10.7717/peerj.542.

69. Mauleón E, Hillán L, Moreno L, Gómez I, Bordons M. Assessing gender balance among journal authors and editorial board members. Scientometrics. 2013;95(1):87–114. doi:10.1007/s11192-012-0824-4.

70. Metz I, Harzing AW. Gender Diversity in Editorial Boards of Management Journals. Academy of Management Learning & Education. 2009;8(4):540–557. doi:10.5465/amle.8.4.zqr540.

71. Metz I, Harzing AW. An update of gender diversity in editorial boards: a longitudinal study of management journals. Personnel Review. 2012;41(3):283–300. doi:10.1108/00483481211212940.

72. Morton MJ, Sonnad SS. Women on professional society and journal editorial boards. Journal of the National Medical Association. 2007;99(7):764–771.

73. Stegmaier M, Palmer B, Assendelft Lv. Getting on the Board: The Presence of Women in Political Science Journal Editorial Positions. PS: Political Science & Politics. 2011;44(4):799–804. doi:10.1017/S1049096511001284.

74. Topaz CM, Sen S. Gender Representation on Journal Editorial Boards in the Mathematical Sciences. PLOS ONE. 2016;11(8):e0161357. doi:10.1371/journal.pone.0161357.

75. Espin J, Palmas S, Carrasco-Rueda F, Riemer K, Allen PE, Berkebile N, et al. A persistent lack of international representation on editorial boards in environmental biology. PLOS Biology. 2017;15(12):e2002760. doi:10.1371/journal.pbio.2002760.

76. Addis E, Villa P. The Editorial Boards of Italian Economics Journals: Women, Gender, and Social Networking. Feminist Economics. 2003;9(1):75–91. doi:10.1080/1354570032000057062.

77. Avin C, Keller B, Lotker Z, Mathieu C, Peleg D, Pignolet YA. Homophily and the Glass Ceiling Effect in Social Networks. In: Proceedings of the 2015 Conference on Innovations in Theoretical Computer Science. ITCS’15. New York, NY, USA: ACM; 2015. p. 41–50. Available from: http://doi.acm.org/10.1145/2688073.2688097.

78. Szell M, Thurner S. How women organize social networks different from men. Scientific Reports. 2013;3:1214. doi:10.1038/srep01214.

79. Lerback J, Hanson B. Journals invite too few women to referee. Nature News. 2017;541(7638):455. doi:10.1038/541455a.

80. Tite L, Schroter S. Why do peer reviewers decline to review? A survey. Journal of Epidemiology & Community Health. 2007;61(1):9–12. doi:10.1136/jech.2006.049817.

81. Bol T, Vaan Md, Rijt Avd. The Matthew effect in science funding. Proceedings of the National Academy of Sciences. 2018; p. 201719557. doi:10.1073/pnas.1719557115.

82. O’Brien KR, Hapgood KP. The academic jungle: ecosystem modelling reveals why women are driven out of research. Oikos. 2012;121(7):999–1004. doi:10.1111/j.1600-0706.2012.20601.x.

83. Petersen AM, Penner O. Inequality and cumulative advantage in science careers: a case study of high-impact journals. EPJ Data Science. 2014;3(1):24. doi:10.1140/epjds/s13688-014-0024-y.

84. Day TE. The big consequences of small biases: A simulation of peer review. Research Policy. 2015;44(6):1266–1270. doi:10.1016/j.respol.2015.01.006.

85. Tomkins A, Zhang M, Heavlin WD. Reviewer bias in single-versus double-blind peer review. Proceedings of the National Academy of Sciences. 2017; p. 201707323. doi:10.1073/pnas.1707323114.

86. Valkonen L, Brooks J. Gender balance in Cortex acceptance rates. Cortex. 2011;47(7):763–770. doi:10.1016/j.cortex.2011.04.004.

87. Fox CW, Burns CS, Meyer JA, Thompson K. Editor and reviewer gender influence the peer review process but not peer review outcomes at an ecology journal. Functional Ecology. 2015;30(1):140–153. doi:10.1111/1365-2435.12529.

88. Borsuk RM, Aarssen LW, Budden AE, Koricheva J, Leimu R, Tregenza T, et al. To Name or Not to Name: The Effect of Changing Author Gender on Peer Review. BioScience. 2009;59(11):985–989. doi:10.1525/bio.2009.59.11.10.

89. Kassis T. How do research faculty in the biosciences evaluate paper authorship criteria? PLOS ONE. 2017;12(8):e0183632. doi:10.1371/journal.pone.0183632.

90. Tregenza T. Gender bias in the refereeing process? Trends in Ecology & Evolution. 2002;17(8):349–350. doi:10.1016/S0169-5347(02)02545-4.

91. Women in neuroscience: a numbers game. Nature Neuroscience. 2006;9(7):853. doi:10.1038/nn0706-853.

92. Bedi G, Van Dam NT, Munafo M. Gender inequality in awarded research grants. Lancet (London, England). 2012;380(9840):474. doi:10.1016/S0140-6736(12)61292-6.

93. Gannon F, Quirk S, Guest S. Searching for discrimination: Are women treated fairly in the EMBO postdoctoral fellowship scheme? EMBO reports. 2001;2(8):655–657. doi:10.1093/embo-reports/kve170.

94. Lee Rvd, Ellemers N. Gender contributes to personal research funding success in The Netherlands. Proceedings of the National Academy of Sciences. 2015;112(40):12349–12353. doi:10.1073/pnas.1510159112.

95. Shen H. Inequality quantified: Mind the gender gap. Nature. 2013;495(7439):22–24. doi:10.1038/495022a.

96. Pohlhaus JR, Jiang H, Wagner RM, Schaffer WT, Pinn VW. Sex Differences in Application, Success, and Funding Rates for Nih Extramural Programs. Academic Medicine. 2011;86(6):759–767. doi:10.1097/ACM.0b013e31821836ff.

97. Waisbren SE, Bowles H, Hasan T, Zou KH, Emans SJ, Goldberg C, et al. Gender differences in research grant applications and funding outcomes for medical school faculty. Journal of Women’s Health (2002). 2008;17(2):207–214. doi:10.1089/jwh.2007.0412.

98. Marsh HW, Jayasinghe UW, Bond NW. Gender differences in peer reviews of grant applications: A substantive-methodological synergy in support of the null hypothesis model. Journal of Informetrics. 2011;5(1):167–180. doi:10.1016/j.joi.2010.10.004.

99. Lariviere V, Sugimoto C, Tsou A, Gingras Y. Team size matters: Collaboration and scientific impact since 1900. arXiv:14108544 [cs]. 2014;.

100. Wuchty S, Jones BF, Uzzi B. The Increasing Dominance of Teams in Production of Knowledge. Science. 2007;doi:10.1126/science.1136099.

101. Larivière V, Desrochers N, Macaluso B, Mongeon P, Paul-Hus A, Sugimoto CR. Contributorship and division of labor in knowledge production. Social Studies of Science. 2016;46(3):417–435. doi:10.1177/0306312716650046.

102. Lloyd ME. Gender factors in reviewer recommendations for manuscript publication. Journal of Applied Behavior Analysis. 1990;23(4):539–543. doi:10.1901/jaba.1990.23-539.

103. Wing DA, Benner RS, Petersen R, Newcomb R, Scott JR. Differences in editorial board reviewer behavior based on gender. Journal of Women’s Health (2002). 2010;19(10):1919–1923. doi:10.1089/jwh.2009.1904.

104. Bornmann L, Daniel HD. Gatekeepers of science—Effects of external reviewers’ attributes on the assessments of fellowship applications. Journal of Informetrics. 2007;1(1):83–91. doi:10.1016/j.joi.2006.09.005.

105. Petty RE, Fleming MA, Fabrigar LR. The Review Process at PSPB: Correlates of Interreviewer Agreement and Manuscript Acceptance. Personality and Social Psychology Bulletin. 1999;25(2):188–203. doi:10.1177/0146167299025002005.

106. Zhang X. Effect of reviewer’s origin on peer review: China vs. non-China. Learned Publishing. 2012;25(4):265–270. doi:10.1087/20120405.

107. Gelman A, Stern H. The Difference Between “Significant” and “Not Significant” is not Itself Statistically Significant. The American Statistician. 2006;60(4):328–331. doi:10.1198/000313006X152649.

108. Adamo SA. Attrition of Women in the Biological Sciences: Workload, Motherhood, and Other Explanations Revisited. BioScience. 2013;63(1):43–48. doi:10.1525/bio.2013.63.1.9.

109. Ceci SJ, Williams WM, Barnett SM. Women’s underrepresentation in science: sociocultural and biological considerations. Psychological Bulletin. 2009;135(2):218–261. doi:10.1037/a0014412.

110. Ceci SJ, Williams WM. Sex Differences in Math-Intensive Fields. Current Directions in Psychological Science. 2010;19(5):275–279. doi:10.1177/0963721410383241.

111. Xie Y, Shauman KA. Sex Differences in Research Productivity: New Evidence about an Old Puzzle. American Sociological Review. 1998;63(6):847–870. doi:10.2307/2657505.

112. Page SE. The Difference: How the Power of Diversity Creates Better Groups, Firms, Schools, and Societies. Princeton: Princeton University Press; 2008.

113. Campbell LG, Mehtani S, Dozier ME, Rinehart J. Gender-Heterogeneous Working Groups Produce Higher Quality Science. PLOS ONE. 2013;8(10):e79147. doi:10.1371/journal.pone.0079147.

114. Science benefits from diversity. Nature. 2018;558(5). doi:10.1038/d41586-018-05326-3.

115. Giordan M, Csikasz-Nagy A, Collings AM, Vaggi F. The effects of an editor serving as one of the reviewers during the peer-review process. F1000Research. 2016;5:683. doi:10.12688/f1000research.8452.1.

116. PEERE policy on data sharing on peer review. PEERE; 2017.

117. Squazzoni F, Grimaldo F, Marušić A. Publishing: Journals could share peer-review data. Nature. 2017;546:352.

118. Nature journals offer double-blind review. Nature News. 2015;518(7539):274. doi:10.1038/518274b.

119. McGillivray B, De Ranieri E. Uptake and outcome of manuscripts in Nature journals by review model and author characteristics. arXiv:180202188 [cs]. 2018;.

120. Ware M. Peer Review in Scholarly Journals: Perspective of the Scholarly Community - Results from an International Study. Inf Serv Use. 2008;28(2):109–112.

121. Kmietowicz Z. Double blind peer reviews are fairer and more objective, say academics. BMJ: British Medical Journal. 2008;336(7638):241. doi:10.1136/bmj.39476.357280.DB.

122. Blank RM. The Effects of Double-Blind versus Single-Blind Reviewing: Experimental Evidence from The American Economic Review. The American Economic Review. 1991;81(5):1041–1067.

123. Okike K, Hug KT, Kocher MS, Leopold SS. Single-blind vs Double-blind Peer Review in the Setting of Author Prestige. JAMA. 2016;316(12):1315–1316. doi:10.1001/jama.2016.11014.

124. Bravo G, Grimaldo F, López-Iñ esta E, Mehmani B, Squazzoni F. The effect of publishing peer review reports on referee behavior in five scholarly journals. Nature Communications. 2019;10(1):322. doi:10.1038/s41467-018-08250-2.

125. Pulverer B. A transparent black box. The EMBO Journal. 2010;29(23):3891–3892. doi:10.1038/emboj.2010.307.

126. Ó Faoleán G. Frontiers Collaborative Peer Review: criteria to accept and reject manuscripts; 2016. Available from: https://blog.frontiersin.org/2016/09/20/the-review-process-making-decisions-on-manuscripts/.

127. Merchant S, Eckardt NA. The Plant Cell Begins Opt-in Publishing of Peer Review Reports. The Plant Cell. 2016;28(10):2343. doi:10.1105/tpc.16.00798.

128. Pourquié O, Brown K. Future developments: your thoughts and our plans. Development (Cambridge, England). 2016;143(1):1–2. doi:10.1242/dev.133355.

129. Rodgers P. Peer Review: Decisions, decisions. eLife. 2017;6:e32011. doi:10.7554/eLife.32011.

130. Abdill, Richard J. and Blekhman, Ran Tracking the popularity and outcomes of all bioRxiv preprints. bioRxiv. 2019; p. 515643 doi:10.1101/515643

